# Pre-existing and emergent cortical neuronal assembly sequences during learning

**DOI:** 10.1101/2025.11.12.688154

**Authors:** Luke Pemberton, Huijeong Jeong, Vijay Mohan K Namboodiri

## Abstract

Neuronal assemblies — groups of co-active neurons — support memory consolidation and retrieval. In the hippocampus, assemblies can pre-exist learning and contribute to memory through sequential activation. Whether similar principles apply to higher cortical areas for flexible memory storage like the orbitofrontal cortex (OFC) remains unclear. Using a novel ground truth-validated clustering approach, we investigated the activity of longitudinally tracked mouse OFC neurons during cue-reward memory acquisition and maintenance. Assemblies active after learning pre-existed the learning and exhibited two distinct sequential dynamics consistent with memory consolidation or retrieval. Consolidation sequences emerged during learning, while retrieval sequences partly recruited pre-existing reward sequences. These findings demonstrate that OFC learning recruits pre-existing networks flexibly repurposed for new associations, revealing circuit motifs that may enable cortical memory storage.

## Introduction

Neuronal assemblies—groups of neurons with high co-activity reflecting shared connectivity—support perception (1–3) and memory (4–8). In the visual cortex, distinct assemblies (also called ensembles) represent different sensory percepts (2, 9, 10), while in the hippocampus, distinct assemblies represent spatial locations (7, 11–13), sensory episodes (14– 17), or components of a cognitive map (18–21). In both sensory cortex and hippocampus, some assemblies pre-exist learning (9, 22–25), though the extent of pre-existence versus emergence during learning remains debated (26–29). Assemblies can also form sequences (9), particularly in the hippocampus (30, 31). For example, the hippocampus replays sequences of assemblies corresponding to past experiences at compressed (32, 33) or experienced time scales (34). Such awake replay is thought to contribute to memory consolidation by repeatedly reactivating prior sensory sequences (35–39). During navigation, compressed sequences can sweep ahead of the animal’s position (theta sequences), possibly reflecting retrieval of upcoming or planned locations (40–43). In sum, assembly sequences may underlie both memory consolidation and retrieval.

Whether assembly sequences that may support memory consolidation and retrieval also exist in higher cortical areas involved in memory storage remains poorly understood. We examined this question in the orbitofrontal cortex (OFC) because similar to the hippocampus, it is important for representing cognitive maps/variables (44–49), values (50– 54), or associations (55–59)—functions that may rely on flexible neuronal assembly sequences. To this end, we developed a ground truth–validated clustering approach for identifying pooled assemblies across animals and further validated it using electrophysiological recordings from neurons throughout the visual cortical hierarchy. We then applied this standardized clustering method to analyze assembly sequence dynamics in OFC neurons longitudinally tracked during cue–reward memory acquisition and maintenance. We found distinct dynamics in the evolution of neuronal assembly sequences consistent with memory consolidation and retrieval.

## Results

### Assembly detection by clustering of event-triggered activity

Identifying neuronal assemblies by assessing co-activity requires simultaneous recordings from neurons over sufficiently long time periods. As a result, commonly used assembly detection methods operate on the activity of neurons recorded simultaneously during a single session in a single behaving animal (60–64). The functional relevance of these assemblies is typically evaluated by analyzing their activity time-locked to behavioral events. This strategy can then be extended to pool functionally comparable assemblies across animals, though this often results in high redundancy, with many assemblies—sometimes more than 20—relating to a single behavior and overlapping with assemblies associated with other behaviors (65).

In contrast, we developed a different approach for assembly detection. We hypothesized that neurons co-active outside of task periods, when shared inputs are minimal, are also likely co-active during task periods when shared inputs are pronounced. Thus, we reasoned that neurons with highly similar peri-event time histograms (PETHs)—reflecting high co-activity around behavioral events—may also be co-active outside those events, thereby forming assemblies. This rationale enables pooling of neurons across animals by evaluating their PETH similarity (**Fig 1A**). After assigning non-overlapping group memberships to neurons with similar PETHs using dimensionality reduction and clustering, we then assess non-event-aligned co-activity among simultaneously recorded neurons. This allows us to determine whether the clusters represent assemblies, specifically by examining whether within-cluster correlations exceed between-cluster correlations (**Fig 1A**). Unlike conventional assembly detection methods, the distinct assemblies detected by this method are guaranteed to have non-redundant task activity.

**Fig. 1.**
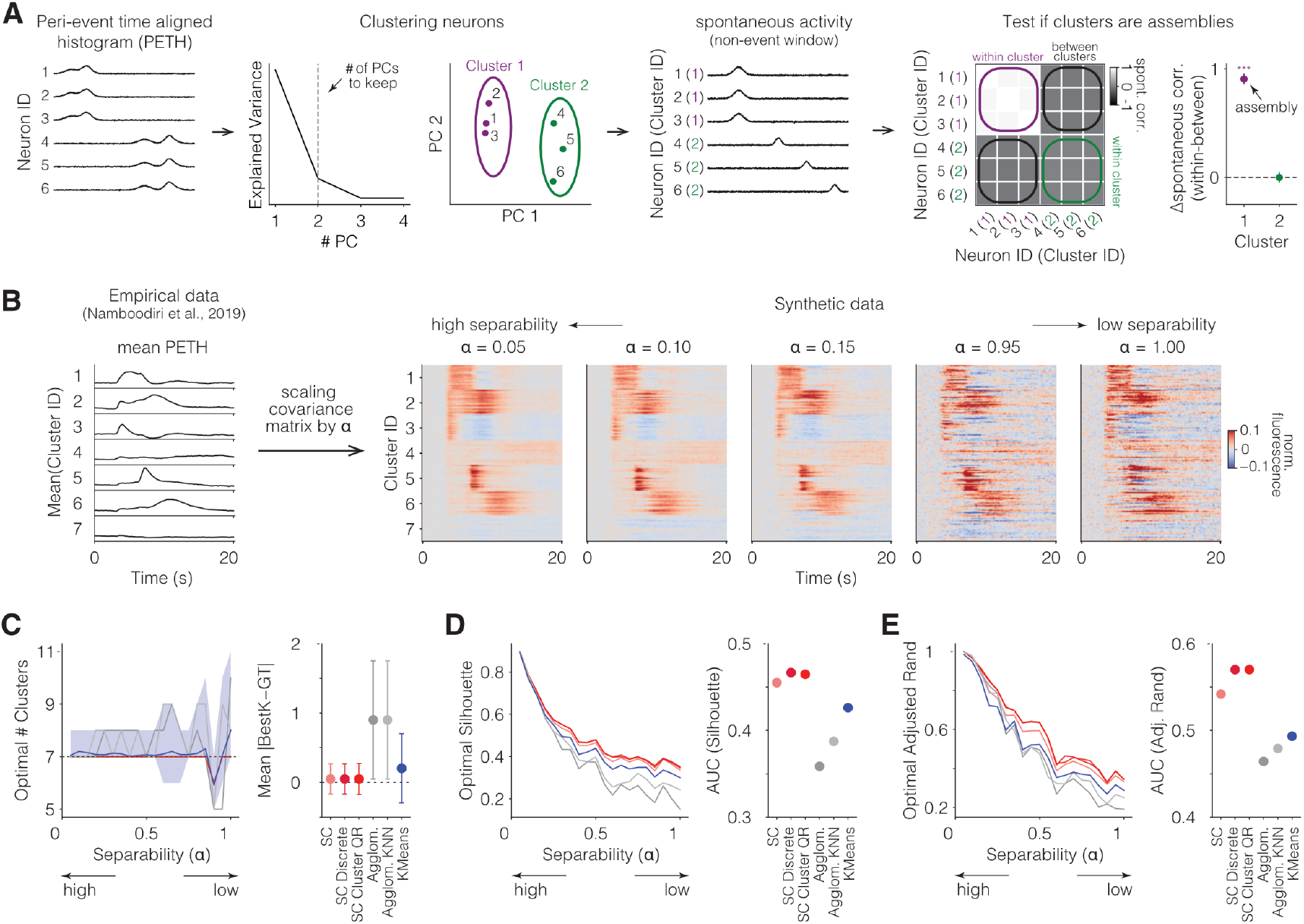
Discrete spectral clustering is the best ground truth validated clustering method for identifying PETH similarity. **A**. Schematic of clustering pipeline for assembly detection involving clustering of task-related PETHs and subsequent testing of whether these clusters are assemblies during non-task aligned activities. Neurons clustered by their PETHs are then tested for co-activity fluctuations during non-event periods. Only clusters with high co-activity (i.e., high within-cluster correlation minus between-cluster correlation) are considered assemblies. See Fig S1 for all clustering algorithms tested here. **B**. First ground truth simulated dataset for testing different clustering algorithms (other datasets in Fig S2, S3). Here, we used a previously published dataset to simulate seven possible PETHs (55). The original covariance matrix across the seven possible clusters was multiplied by the scale factor *α*. Low values of *α* increase cluster separability by retaining mean locations in activity space while reducing variance. 100 neurons with each PETH type were simulated with random Gaussian noise added to each neuron’s PETH (Methods). *α* = 1 represents separability comparable to the original data (but in a ground truth simulation). **C**. Identified number of clusters based on six different clustering algorithms to separate neurons by their PETH types. Seven is the ground truth number of clusters. Different algorithms are represented by different colors (spectral clustering variants are a similar shade), as shown in the plot on the right measuring the deviation between the identified number of clusters (“BestK”) and the ground truth number of clusters (“GT”). **D**. The silhouette score, a measure of cluster separation, for the identified optimal number of clusters for each algorithm. **E**. The adjusted rand index, a measure of how correctly each neuron was assigned to its ground truth cluster, for each algorithm. Across the different metrics shown in C-E, discrete spectral clustering performs the best.

To identify the best clustering algorithm for detecting PETH similarity, we simulated realistic ground truth data of clusters with many distinct PETH shapes, and parametrically varied cluster separability (Methods). We tested many common clustering methods, some shown in **Fig 1B** and others in **Fig S1**. When ground truth clusters are well separated, all methods performed equally well. However, at reduced separability, discrete spectral clustering (66) proved to be the most effective (**Fig 1C-E**). This approach was also the best on less realistic simulated ground truth data of different types (**Figs S2, S3**). Across these ground truth datasets, the benefit of using discrete spectral clustering was higher when the simulated PETHs differed in shape (**Figs 1, S2**) and not just the location of similarly shaped peak activity (**Fig S3**). Together, our findings indicate that discrete spectral clustering is the best method for identifying neurons with high PETH co-activity.

We next tested whether the clusters identified using this method are assemblies using a large-scale single unit electrophysiology dataset across the visual cortical hierarchy (67), similar to what has been reported (68) (**Fig S4**). Instead of aligning activity on an external behavioral event, here we aligned the activity of neurons to hippocampal ripples, an internal brain event. We identified three neuronal clusters based on ripple-aligned PETHs. The activity of these clusters showed higher within-cluster co-activity outside of ripple periods (**Fig S4**), validating that identified clusters are indeed assemblies. Interestingly, the sequential activity of these assemblies was conserved across both ripple and non-ripple periods, proceeding in a non-time compressed manner. Thus, event-aligned co-activity using discrete spectral clustering can indeed identify true neuronal assemblies and enable investigation of assembly sequences.

### OFC neuronal assemblies detected after cue-reward learning pre-exist the learning

We next applied this method to identify neuronal assemblies while animals experienced sequential pairings of a cue followed by reward (i.e., cue-reward learning) in the ventral/medial OFC. The OFC is anatomically and functionally heterogeneous, with medial (MO), ventral (VO), and lateral (LO) subregions differing in proposed roles in value and task-state representation (47, 54, 58). We focused on ventral/medial OFC because prior work implicates this region in the encoding of long-term cue–outcome memories (55, 69), making it a strong candidate for assembly-based memory sequences.

Briefly, head-fixed mice learned the association between an auditory tone lasting 2s (CS+) and a reward after a 1s gap (“trace interval”); another auditory tone (CS-) was not paired with reward (**Fig 2A**). Trials were followed by variable intertrial intervals (ITI; constant 3s plus an exponential distribution of mean 30s). Mice learned this association in a week on average. After stable behavior was reached (referred to as “trained day”), two forms of contingency degradation were performed in a random order (either reduction of reward probability to 50%, or unpredicted rewards in the inter-trial interval, see Methods). After retraining to full contingency, mice finally underwent extinction of the cue-reward pairing, i.e., reward probability was reduced to zero. To assess neuronal encoding during initial learning, we focused on the early days of learning until the trained day. To assess memory maintenance, we compared neuronal encoding between the trained day and the days before and after extinction. Conditioned behavior, as measured by anticipatory licking, was low on early days and high after learning as well as the day before extinction, and low after extinction (55) (**Fig 2B**). OFC neurons were targeted using an AAV expressing GCaMP6s under the CaMKIIα promoter (**Fig 2C**). Because cue responses may reflect sensory processing and/or reward association, while emergent trace interval responses only reflect cue-reward association due to lack of sensory input, we first assessed “neural learning” by measuring the evolution of trace interval responses. Average neuropil corrected CS+ trace interval fluorescence across the entire longitudinally tracked neuronal population (n=1,435 neurons from n=5 mice) was low on early days of learning, high after learning and low after extinction (**Fig 2D**). Collectively, these results demonstrate that animals did not exhibit evidence of behavioral or neural learning on day 1 of conditioning, allowing a timepoint to evaluate pre-existing network structure.

**Fig. 2.**
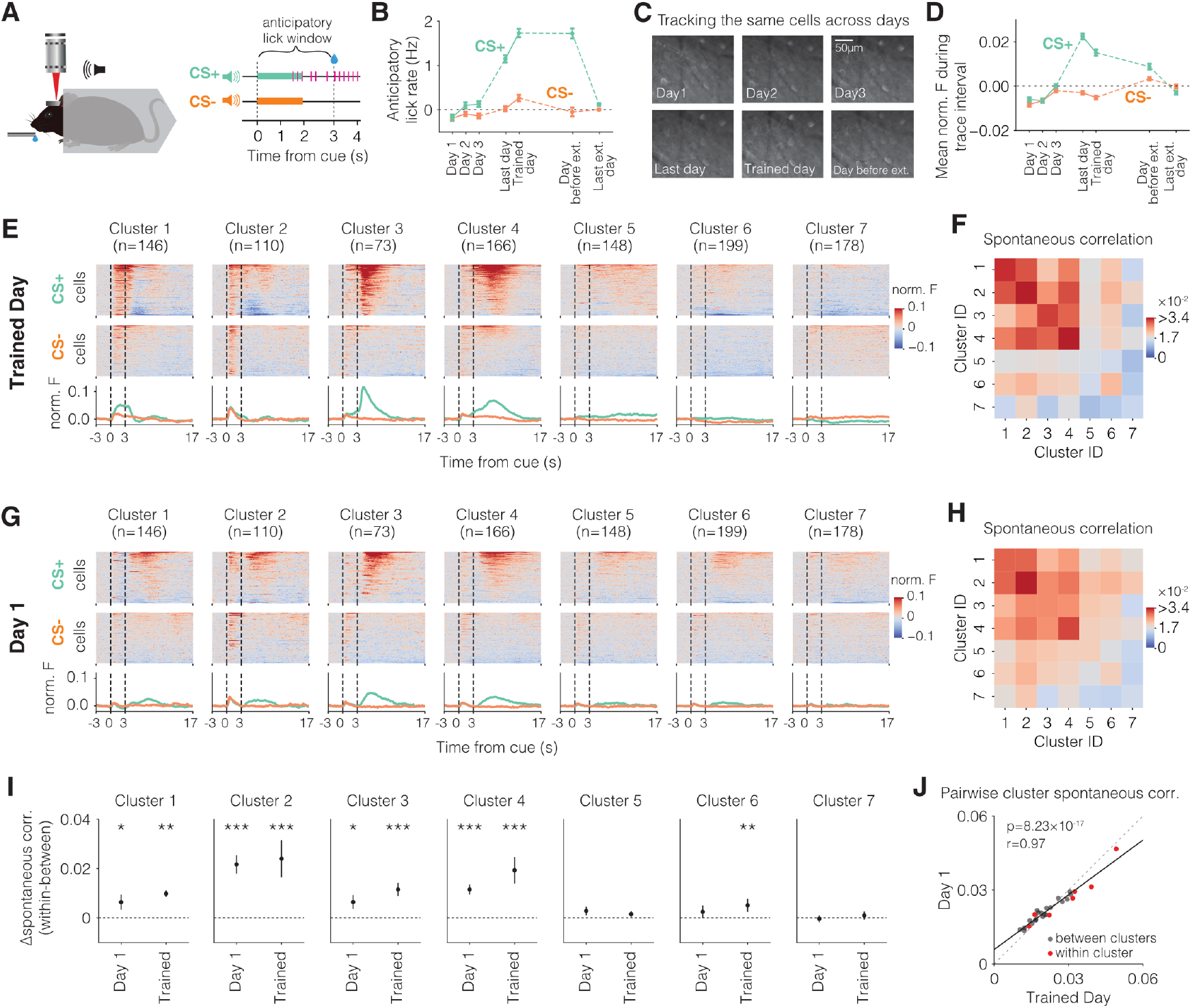
OFC neuronal assemblies identified after learning pre-exist learning. **A**. Schematic of head-fixed two-photon imaging and trace conditioning paradigm. Anticipatory lick rate is measured during the cue-outcome period to assess behavioral evidence reflecting cue-reward memory. **B**. Baseline subtracted anticipatory lick rate across various task days, demonstrating clear behavioral evidence of acquired cue-reward memory on the last day of learning (i.e., day before trained day), trained day, and day before extinction. Baseline period was 3 s before cue. Data from n=5 mice. During extinction, reward was not delivered after CS+, which eliminates anticipatory licking. Last day of learning was day 7.2 on average across mice (median: 5 range:[5, 11]). Trained day was day 8.2 on average across mice (median: 6 range:[6, 12]). Day before extinction was day 21.8 on average across mice (median: 21 range:[18, 26]). Last day of extinction was day 24.4 on average across mice (median: 24 range:[22, 28]) (see Methods for detailed timeline of experiments). Error bars are s.e.m. **C**. Example cropped field of view showing longitudinal tracking of the same neurons across days. **D**. Mean normalized fluorescence (see Methods) during the trace interval (2-3 s after CS+ onset) across all longitudinally tracked neurons (n=1,435 neurons across 5 mice), showing that there is no evidence of neural learning on day 1. Error bars are s.e.m. **E**. Results of discrete spectral clustering applied to OFC neurons based on their CS+ and CS-evoked PETHs. Only neurons that were longitudinally tracked throughout learning are included. Top two rows show CS+ and CS-aligned activity of all neurons within each cluster. The neurons are sorted by their total activity during CS+ trials and arranged in the same order for CS-. **F**. Heatmap showing spontaneous correlations between each pair of clusters based on their inter-trial interval (ITI) activity outside of CS+ or CS-trials. **G**. Same as E but for day 1 of conditioning. **H**. Same as F but measured using ITIs on day 1 of conditioning. **I**. The difference in within-cluster versus between-clusters spontaneous correlations during the ITI on both day 1 and trained day. Clusters 1-4 are assemblies on both days. *p<0.05, **p<0.01, ***p<0.001. Error bars are s.e.m. estimated from a hierarchical bootstrap. **J**. Scatter plot of the spontaneous correlations between each pair of clusters measured on day 1 and trained day. Within-cluster and between-clusters pairs are separated. *<u>See Table 1 for all statistical results and details</u>*.

We used discrete spectral clustering to cluster neurons based on their PETHs on CS+ and CS-trials after removing non-responsive neurons (Methods). We identified seven neuronal clusters with distinct PETHs (**Fig 2E**). This contrasts with a conventional assembly detection algorithm, which exhibited highly redundant task activity (**Fig S5**). The activity of these clusters on day 1 did not differ between CS+ and CS-trials prior to the outcome period (**Fig 2F**). We next assessed whether these identified clusters were assemblies by measuring within-cluster vs between-cluster correlations during the ITI. When measuring ITI correlations, we used deconvolved fluorescence signals that have been shown to approximate spiking using ground truth ex vivo OFC recordings (Supplementary Fig 1 in (55)) and using a meta-analysis of deconvolution methods (70). On the trained day, clusters 1-4 and 6 showed higher within-cluster than between-cluster correlation, indicating that they are assemblies (**Fig 2G**, all p values were obtained by hierarchical bootstrapping (71) and corrected for multiple comparisons (see Methods and **Fig S6**). Clusters 1-4 also showed higher within-cluster than between-cluster correlation on day 1, indicating that the assemblies pre-existed cue-reward learning (**Fig 2G**). We next plotted the average ITI correlation between simultaneously recorded neurons between all cluster pairs and found that this overall network correlation structure looked remarkably similar between day 1 and trained days (**Fig 2H**). Indeed, the correlation between ITI correlations of every cluster pair across the two days was extremely high, with a slope less than 1 indicating that the network correlation structure strengthened the higher within-cluster correlations and weakened the lower between-cluster correlations (**Fig 2H**). In sum, OFC neuronal assemblies identified based on their PETHs on the trained day pre-existed and strengthened with learning.

### OFC neuronal assembly sequences consistent with memory consolidation and retrieval

While memory retrieval is triggered by the cue, memory consolidation, by definition, requires cue-reward memory representations to be activated outside the cue-reward experiences (e.g., like hippocampal replay during rest (35)). Because animals sequentially experienced the cue and reward on individual CS+ trials, we reasoned that assemblies encoding the cue-reward memory may form sequences. Therefore, we next investigated whether there are any sequential dynamics of the OFC neuronal assemblies consistent with memory consolidation and retrieval. Here, we focused on the four main clusters with strong cue and reward evoked responses: 1, 2, 3, and 4 (**Fig 3A**). Clusters 1 and 2 strongly respond to the cue, while clusters 3 and 4 strongly respond to reward. Visualization of the CS+ PETHs suggested a sequence of activity among these clusters from 2 to 1 to 3 to 4. However, this apparent sequence on CS+ trials may just be an artifact of the fact that clusters 1 and 2 mainly respond to cues, and 3 and 4 mainly respond to rewards. Indeed, the ordering between reward responses for clusters 3 and 4 appears to be maintained on day 1 with cluster 3 exhibiting short latency reward responses and cluster 4 exhibiting long latency reward responses (**Fig 2F**).

**Fig. 3.**
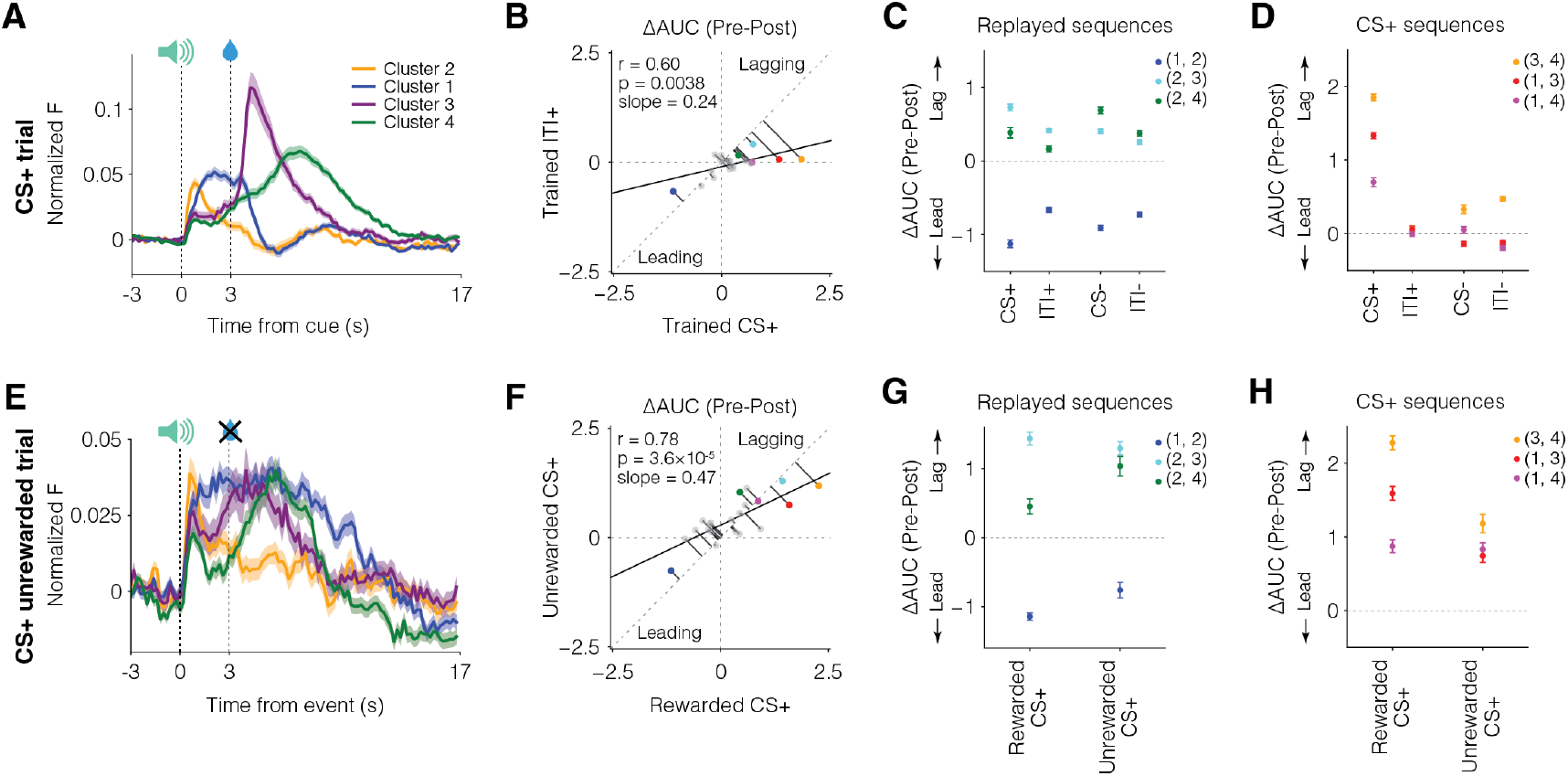
OFC assembly sequence dynamics after cue-reward learning are consistent with memory consolidation and retrieval. **A**. CS+ aligned PETHs for clusters 1-4 on the trained day. The sequence 2 → 1 → 3 → 4 is visible. Error shading is s.e.m. **B**. The sequential relationship between assemblies was measured using the area under the curve of the cross correlogram (CCG) over negative lags minus the corresponding area over positive lags (referred to as ΔAUC) (**Fig S7**). Here, ΔAUC is plotted for all cluster pairs during CS+ periods (0-17 s following CS+ onset) and ITI+ periods (ITIs following CS+ trials). Weak but significant correlation in these ΔAUCs implies that some sequences are replayed during ITI while others are not. **C**. ΔAUC of “replayed sequences” during CS+, ITI+, CS- and ITI-periods, showing maintenance of ΔAUCs across all time periods. Note that all replayed sequences include cluster 2. Error bars are 95% confidence intervals estimated from a hierarchical bootstrap. **D**. ΔAUC of “CS+ sequences” during CS+, ITI+, CS- and ITI-periods, showing much larger ΔAUCs during CS+ compared to other time periods. Note that no CS+ sequence includes cluster 2. Error bars are 95% confidence intervals estimated from a hierarchical bootstrap. **E**. CS+ aligned PETHs on unrewarded trials during a 50% reward probability session after the trained day (see Methods). The sequence 2 → 1 → 3 → 4 is still visible. Error shading is s.e.m. **F**. ΔAUC is plotted for all cluster pairs during CS+ rewarded vs unrewarded periods on the 50% reward probability session. **G**. ΔAUC of “replayed sequences” during CS+ rewarded and unrewarded periods, showing that all sequences are present on unrewarded trials. Error bars are 95% confidence intervals estimated from a hierarchical bootstrap. **H**. ΔAUC of “CS+ sequences” during CS+ rewarded and unrewarded periods, showing that all sequences are present on unrewarded trials. Error bars are 95% confidence intervals estimated from a hierarchical bootstrap.

During the cue-reward experience itself, sequential activation of assemblies might simply reflect the sequential experience of cue and reward. To rule out trivial sequences observed only during the experimentally induced cue-reward pairing, we first assessed sequences during the ITI. We measured sequences between a pair of assemblies during the CS+ trials and the ITIs following CS+ trials (ITI+) using cross-correlation analysis. We identified the asymmetry in the cross-correlogram by measuring the difference between the negative lag area under the curve and the positive lag area under the curve (ΔAUC) (**Fig S3**). Zero ΔAUC indicates no sequential relationship between the pair of clusters and high ΔAUC suggests strong sequential relationship. We plotted the correlation between ΔAUC for every pair of clusters (including 5-7) between CS+ and ITI+ periods on the trained day and found a significant, but weak correlation of slope less than 1 (**Fig 3B**). This suggests that some cluster pairs exhibited similar sequential relationships between CS+ and ITI+ (consistent with replay), while others exhibited sequential relationships only during CS+ trials. We identified that the assembly pairs (1,2), (2,3) and (2,4) exhibited similar sequences between CS+, ITI+, CS-, and ITI-periods (ITI-indicates ITIs following CS-trials). We refer to these as replayed sequences (**Fig 3C**). We also measured the centroid lag of the cross-correlogram (**Fig S7**) and found that the lag between these sequences was similar, but slightly smaller, between CS+ periods with cue-reward pairing and the other periods, suggesting that the CS+-reward memory may be replayed by these sequences at approximately the experienced cue-reward lag, consistent with a memory consolidation mechanism (**Fig 3C**).

On the other hand, the (1,3), (1,4) and (3,4) sequences were much stronger in the CS+ periods compared to CS-, ITI+, or ITI-periods. We refer to these as CS+ sequences (**Fig 3D**). The CS+ sequences could trivially be present only during the experimentally induced cue-reward pairing. To rule this out, we measured ΔAUC on CS+ unrewarded trials in the 50% reward probability session (**Fig 3E**). All these sequences maintained ΔAUC on CS+ unrewarded trials (**Fig 3F-H**). The maintenance of the same sequences on CS+ unrewarded trials is consistent with cue-reward memory retrieval.

We next assessed sequence dynamics during and after cue-reward learning (**Figs 4, S8**). We discovered that replayed sequences emerged during learning, i.e., were weak on days 1-3, and were thereafter maintained on CS+, ITI+, CS-, and ITI-periods even after extinction (**Fig 4A-D**). Thus, these sequences are consistent with a mechanism that consolidates an acquired cue-reward memory. The fact that these sequences were maintained even after extinction when the conditioned behavior was abolished demonstrates that their activation is not a pure reflection of licking behavior.

**Fig. 4.**
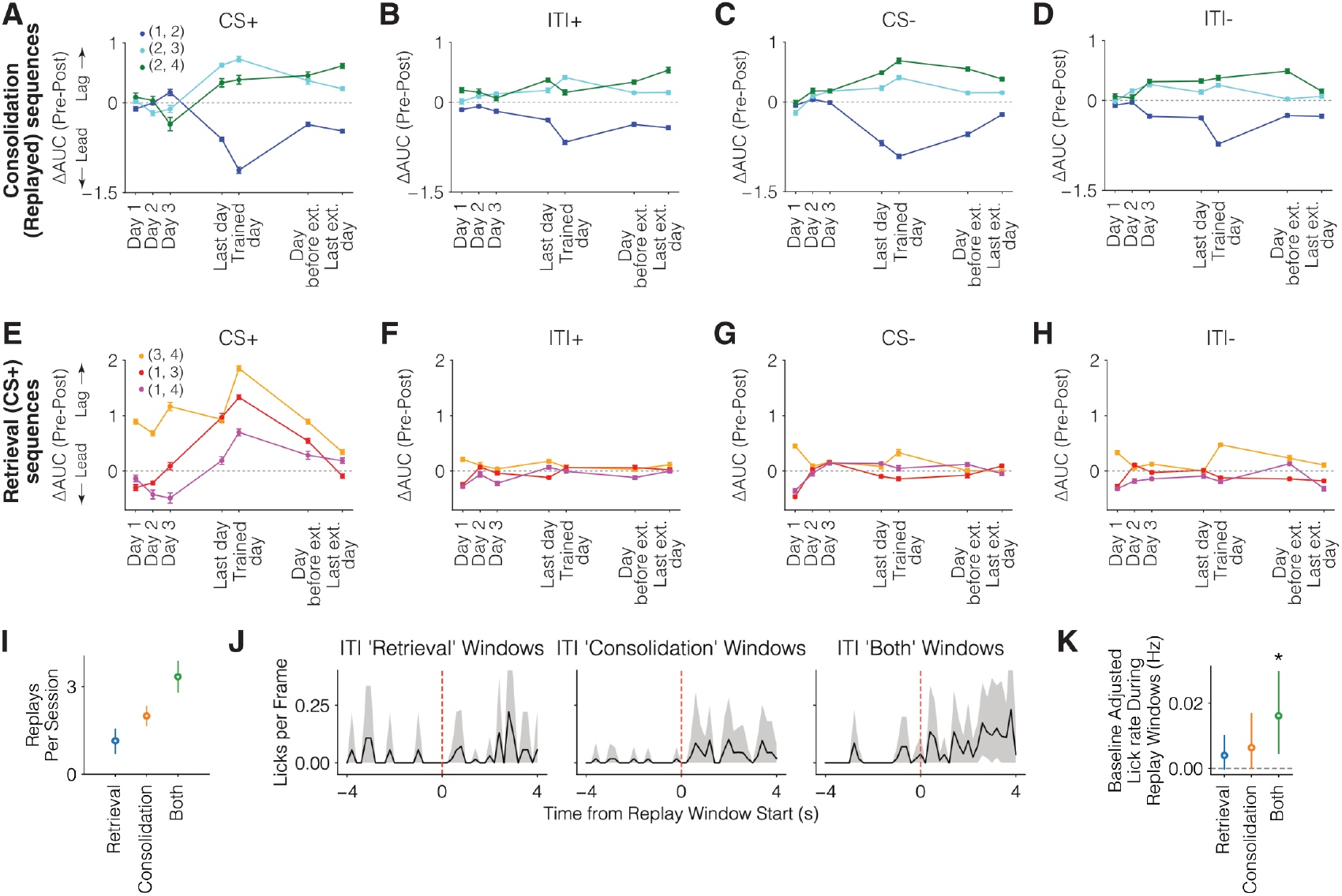
Pre-existing and emergent OFC assembly sequences during cue-reward learning. **A-D**. ΔAUC of replayed sequences across days of interest for all time periods showing emergence during learning and subsequent maintenance even despite behavioral extinction. Error bars are 95% confidence intervals estimated from a hierarchical bootstrap. **E-H**. ΔAUC of CS+ sequences across days of interest for all time periods showing selective enhancement during CS+ but not other time periods. (3,4) sequence pre-exists learning. While (1,3) and (1,4) also pre-exist learning (ΔAUC<0), their sequential order flips after learning (ΔAUC>0). Error bars are 95% confidence intervals estimated from a hierarchical bootstrap. **I**. Number of identified ITI replays per session of either retrieval only sequences (CS+ sequences), single consolidation pairs (“consolidation” representing a single pair of replayed sequences (1,2), (2,3) or (2,4)), and multiple consolidation pairs, which include both consolidation and retrieval sequences (“both”). To identify these replay windows, all ITI periods were pooled across the last day of learning, trained day, and the day before extinction, i.e., the sessions after learning in which the cue predicted reward with 100% reward probability (see Methods). **J**. PETH of licking aligned to the above replay types. Error shading represents 95% confidence intervals from hierarchical bootstrapping. **K**. Baseline subtracted lick rates during the 4s after replay onset. Error bars represent 95% confidence intervals from hierarchical bootstrapping (Bonferroni corrected, *p<0.05).

Among the CS+ sequences, (1,3), and (1,4) sequences emerged during learning on CS+ trials, but the (3,4) sequence pre-existed learning and was stably maintained during and after learning (**Fig 4E-H**). Interestingly, though the (3,4) sequence is visible after reward on day 1 (**Fig 2F**), it is co-opted as a memory sequence after learning, as evident from its presence on CS+ unrewarded trials (**Fig 3E**) and after cue-reward behavioral extinction (**Fig 4E**). Thus, sequences consistent with memory retrieval partly repurpose a pre-existing reward sequence. Interestingly, consolidation vs retrieval sequences were cleanly separated by the presence or absence, respectively, of cluster 2—the cluster that responds similarly to both CS+ and CS-(**Fig 2E**). This suggests that a generic cue encoding assembly forms learned sequences with other reward-related assemblies to consolidate cue-reward memory.

The above analyses used ΔAUC of cross-correlograms to assess sequential activity. This allows the use of the entire ITI activity to evaluate pairwise lead-lag relationships without identifying individual instances of replay. We next attempted to identify individual instances of replay of every assembly pair of interest (assemblies 1-4) to test whether they had any behavioral correlates. We did so by combining all ITI periods across the last day of learning, trained day, and the day before extinction, i.e., the sessions after learning in which the cue predicted reward with 100% reward probability. To prevent baseline contamination, we identified moments of replay where neither the leading nor lagging assembly was active for 6 seconds prior to the replay onset (which is highly likely to undercount the true number of replays). We classified replays into three categories based on the identity and number of assemblies that constitute each replay: retrieval only, single consolidation pair, or multiple consolidation pairs. The multiple consolidation pairs include a combination of (2,1), (2,3), or (2,4) sequences, and as such, also include sequences of 1, 3, or 4 assemblies (i.e., the retrieval pairs). Therefore, we consider these as both retrieval and consolidation replays. As expected from the earlier cross-correlation analyses, retrieval only replays were rare, while the consolidation replays were more common (**Fig 4I**). The rate of licking was slightly, but significantly higher than baseline after retrieval-consolidation replays (**Fig 4J-K, Fig S9**), suggesting that OFC memory sequence replays during ITI may guide or be guided by licking behavior (though note that consolidation sequences are preserved after behavior extinction). Importantly, because hippocampal ripples and replays are known to occur during the absence of licking (72, 73), these results suggest that OFC assembly replays and hippocampal replays likely do not overlap. Collectively, these results demonstrate that OFC assembly sequences relate to both memory consolidation and retrieval, which are distinguished by the participation or absence of a generic cue encoding assembly (assembly 2).

## Discussion

Cue-reward memories are thought to be encoded in higher-order cortical regions like the OFC by the strength of neuronal activation (55, 56, 74, 75). Here, we show that OFC also repurposes pre-existing neuronal assemblies into sequences consistent with memory retrieval—i.e., induced by the reward predicting cue—and memory consolidation—i.e., induced both by the cue and replayed at moments without the cue. Most of these sequences emerged during learning and were maintained thereafter, although one memory retrieval sequence utilized a pre-existing reward sequence.

Unlike conventional methods, we identified comparable assemblies across animals via task-aligned activities, enabling natural pooling of similar assemblies. Discrete spectral clustering proved optimal for this approach, as task-defined clustering after learning revealed neuronal groups that correspond to pre-existing assemblies during non-task periods before and after learning. A potential limitation is that this method only detects non-overlapping assemblies, while traditional techniques can reveal overlapping membership. However, interpreting task-relevant activity remains a challenge for conventional assembly detection due to highly redundant task-aligned assemblies (**Fig S5**).

The OFC is known to build and represent schema—neural representations of shared rules across disparate tasks (76, 77). Forming schema representations, by definition, requires maintenance of learned memories from one task to subsequently experienced tasks. As such, it may be possible that the pre-existing neuronal assemblies and sequences identified here may provide the neuronal circuit scaffold for schema formation. This is further consistent with the fact that consolidation sequences acquired during cue-reward conditioning are stably maintained despite many task variations including complete behavioral extinction.

Our findings directly relate to two key debates in neuroscience. The first concerns representational drift versus stability (78–83). While representational drift has been reported in several regions even during identical behavioral tasks, this study finds largely stable assemblies and learned sequences persisting across diverse behavioral contingency changes over days. Even after probability or contingency degradations and extinction, many learned sequences remain stable: (1,2), (2,3), (2,4), and (3,4) (**Fig 4**). This stability aligns with recent findings of persistent assemblies in V1 during passive visual exposure (84), and extends them by demonstrating that OFC assembly sequences are stable despite multiple interceding task changes.

The second debate addresses categorical versus non-categorical neural coding. Categorical coding is defined as the organization of individual neurons into functionally distinct subpopulations with distinct neural coding of task variables. Although OFC can exhibit neuronal clusters (55, 85–87), various studies emphasize the prevalence and computational advantage of non-categorical coding without distinct clusters (88–90). Indeed, a recent large-scale survey suggests that categorical coding is uncommon outside sensory cortex (91). Notably, however, the presence of clusters in this study is analyzed using time-averaged activity across many task conditions. Here, in contrast, we show that clustering based on PETHs reliably identifies neuronal clusters that even pre-exist learning and form assembly sequences. We therefore propose temporal activity profiles as critical when assessing the presence of activity-based neuronal clusters. This is consistent with reports of neuronal clusters in many regions identified via PETHs aligned to internal or external events (55, 68, 92–96). In sum, neuronal assemblies with co-active responses may represent a general organizational principle throughout the brain, even if their time-averaged task activity often appears non-categorical.

How are cue-outcome memories stored in the brain? The engram hypothesis, formulated mainly in hippocampus and amygdala, posits that a set of excitable neurons activated during initial cue-outcome experience are captured into a memory representation that can later be retrieved by reactivating the same cells (97). Here, we observed more complex dynamics: although assemblies themselves were present on the first day, the assembly sequences associated with memory consolidation and retrieval largely emerged only after learning, being absent on the first day of experience. This may reflect systems consolidation where memories recruit hippocampus during the early phase of learning but recruit cortex only later (98). Notably, a pre-existing reward sequence present on day 1 later became incorporated into a memory retrieval sequence, suggesting that some sequences present initially are relevant for future memory retrieval.

Cortical assembly sequences identified here, as in the hippocampus, are consistent with a role in memory consolidation and retrieval. However, current technology does not allow direct causal tests of the function of individual assembly sequences in memory. Demonstrating causality would require tools capable of expressing distinct opsins in activity-defined assemblies with unknown genetic or circuit markers—something that remains technically infeasible. The correspondence between activity-defined assemblies and genetically or circuit-defined cell types is unclear, with some studies suggesting a weak link (55, 99). As a result, existing genetic or viral methods cannot label the subsecond-timescale, activity-defined assemblies described here. Further, even though holographic optogenetic stimulation has been used to trigger individual activity-defined assemblies (2, 6), playing back or selectively removing assembly sequences at scale to assess behavioral effects remains technically challenging. Major technological advances will therefore be necessary to replay or remove arbitrary neuronal assembly sequences to directly assess their function. Despite these technical limitations, the present findings demonstrate that the OFC flexibly repurposes pre-existing neuronal assemblies into sequences that may support cue-reward memory consolidation and retrieval.

## ACKNOWLEDGEMENTS

We thank L. Frank, S. Mihalas, M. Kheirbek, J. Berke, and members of the Namboodiri laboratory for helpful discussions. This project was supported by NIH R01MH129582, R01AA029661, the Scott Alan Myers Endowed Professorship, Alfred P Sloan Fellowship, Pew Biomedical Scholarship, and Klingenstein-Simons Fellowship to V.M.K.N. We thank Garret D Stuber in whose lab V.M.K.N. collected OFC imaging data.

## AUTHOR CONTRIBUTIONS

Conceptualization: LP, HJ, VMKN; Methodology: LP, HJ, VMKN; Investigation: LP; Visualization: LP, HJ, VMKN; Funding acquisition: VMKN; Project administration: LP, HJ, VMKN; Supervision: VMKN; Writing – original draft: LP, HJ, VMKN; Writing – review & editing: LP, HJ, VMKN.

## Supplementary Figures

**Fig. S1.**
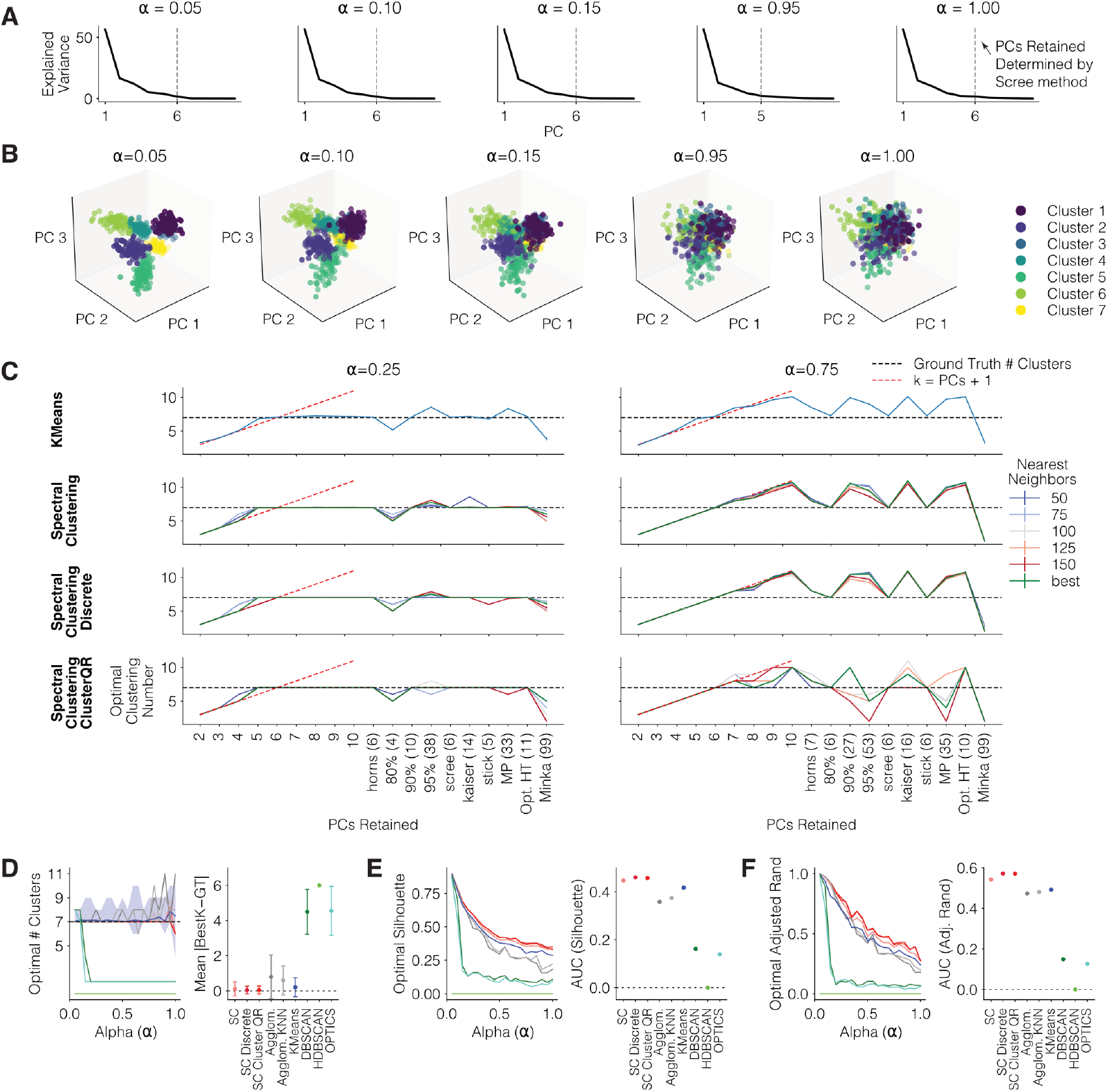
Additional characterization of different clustering algorithms. **A**. Scree plots of the synthetic ground truth datasets shown in **Fig 1** for select values of alpha with the demarcated elbow of the scree curve. **B**. Plots of the same synthetic datasets as in A in 3-dimensional principal component space, showing that clusters are less separable as alpha increases. **C**. The optimal number of clusters recovered from different algorithms using silhouette score as a function of the number of retained principal components (PCs) for two values of alpha. The number of retained PCs determines the identified optimal number of clusters. For low alpha, when the number of retained PCs is too low, the number of clusters identified is lower than the ground truth. Once the number of retained PCs exceeds a threshold, the correct number of clusters is identified. For high alpha, the number of clusters identified exceeds the ground truth when the number of retained PCs keeps increasing. Hence, choosing the correct number of retained PCs is crucial. This relationship was empirically observed for all values of alpha. We empirically observed that finding the scree elbow recovered the ground truth number of PCs. Hence, we recommend this method. By contrast, common approaches such as retaining 80% or 90% total explained variance can result in over or under clustering. **D**. Identified number of clusters based on nine different clustering algorithms to separate simulated neurons by their PETH types. Seven is the ground truth number of clusters. Different algorithms are represented by different colors, as shown in the plot on the right measuring the deviation between the identified number of clusters (“BestK”) and the ground truth number of clusters (“GT”). **E**. The silhouette score, a measure of cluster separation, for the identified optimal number of clusters for each algorithm. **F**. The adjusted rand index, a measure of how correctly each simulated neuron was assigned to its ground truth cluster, for each algorithm.

**Fig. S2.**
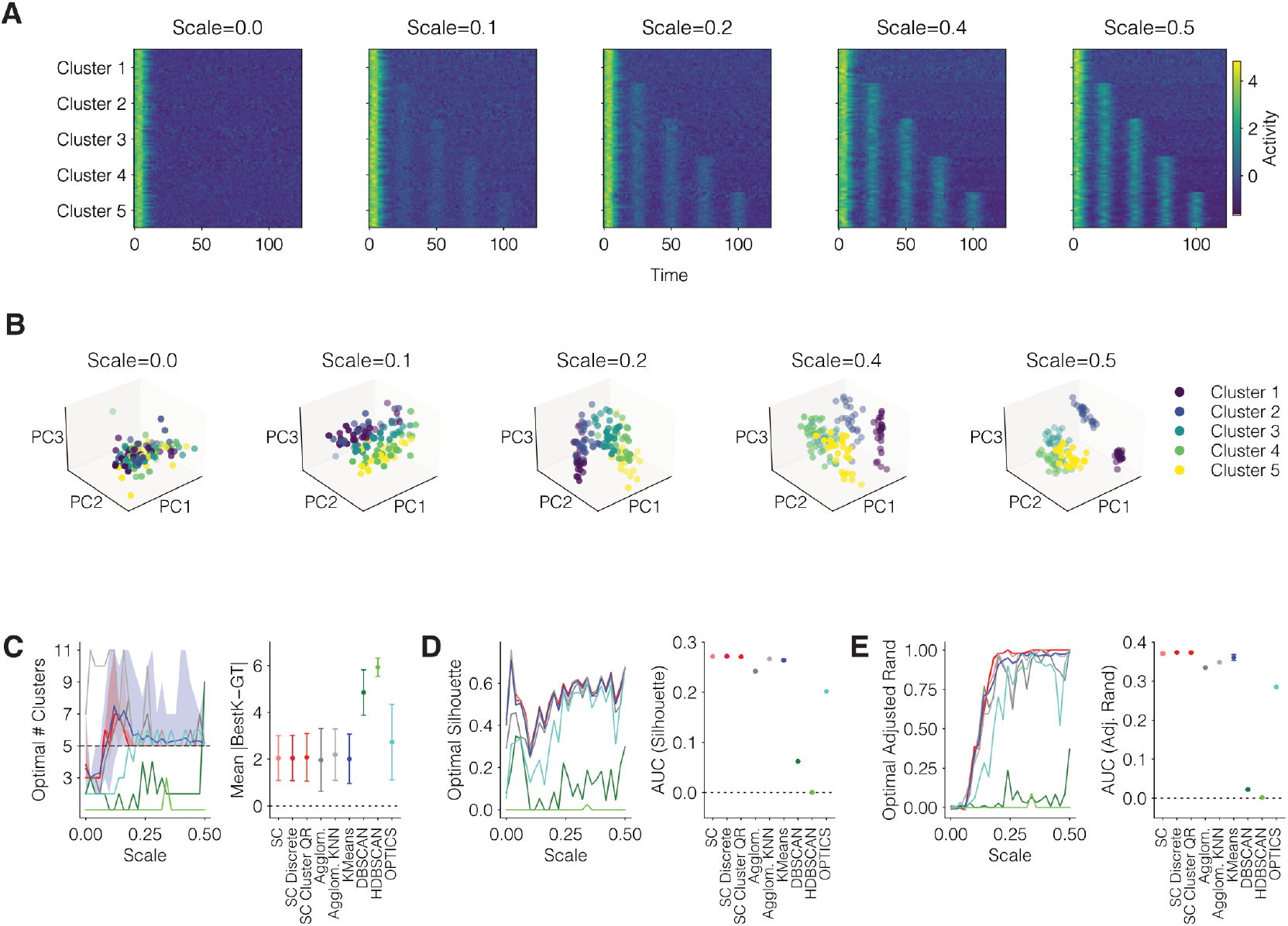
Clustering results on ground truth “Shape” data. **A**. Heatmaps of the “Shape” simulated dataset for select values of the scale dataset parameter. The scale parameter controls the magnitude of the response for 4 additional peaks of activity beyond the first peak at 0. Scale=0 has only one cluster. For other scales, there are 5 ground truth clusters. The clustering becomes easier as the scale parameter increases. **B**. Plots of the same select “Shape” datasets in principal component space. Separability of clusters increases as the scale parameter increases. **C**. Identified number of clusters based on nine different clustering algorithms to separate neurons by their PETH types. Five is the ground truth number of clusters. Different algorithms are represented by different colors, as shown in the plot on the right measuring the deviation between the identified number of clusters (“BestK”) and the ground truth number of clusters (“GT”). **D**. The silhouette score, a measure of cluster separation, for the identified optimal number of clusters for each algorithm. **E**. The adjusted rand index, a measure of how correctly each neuron was assigned to its ground truth cluster, for each algorithm.

**Fig. S3.**
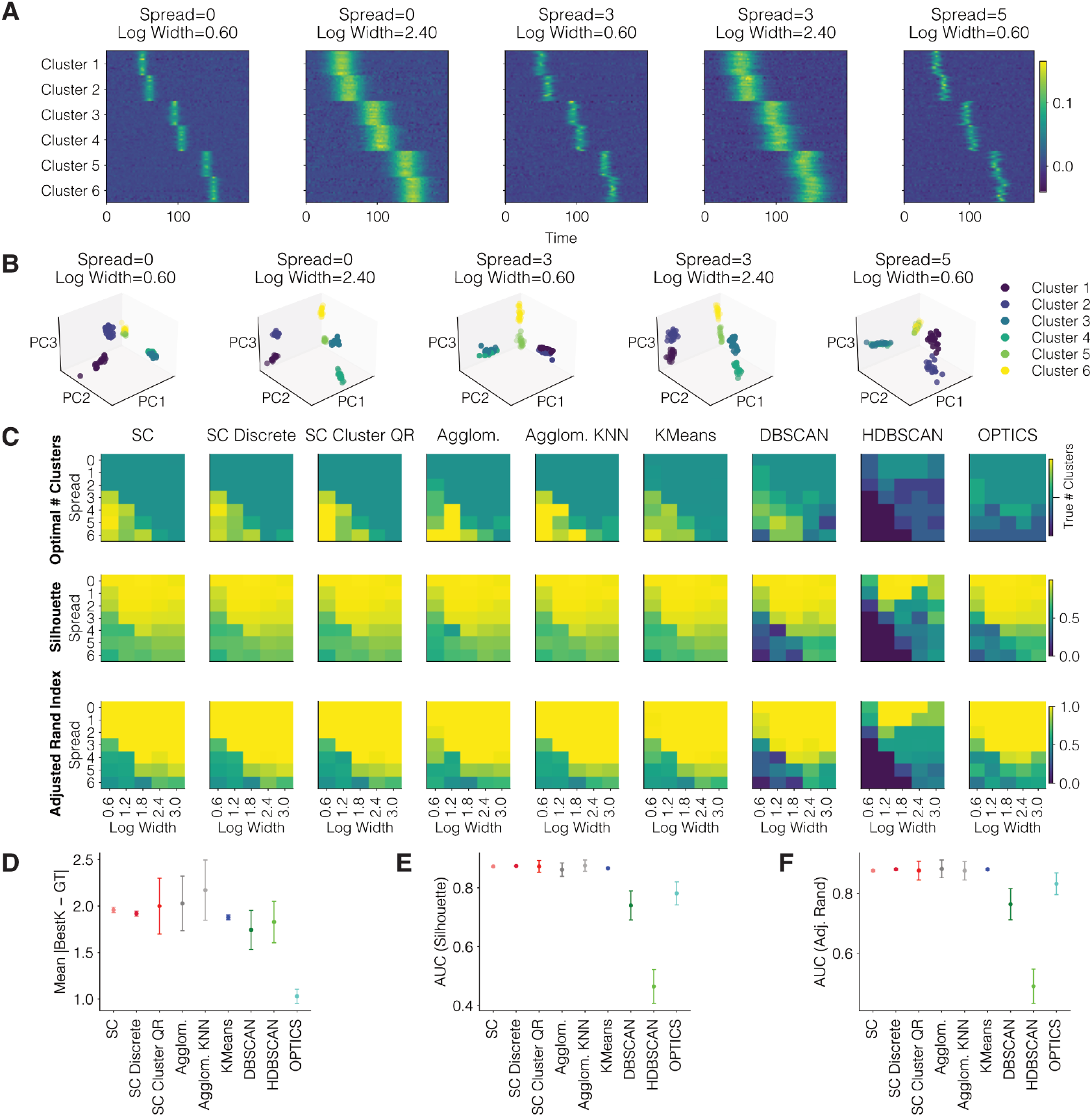
Clustering results on ground truth “Peaks” data. **A**. Heatmaps of the “Peaks” simulated dataset for select values of the spread and width dataset parameters, which together control cluster separability (see Methods). **B**. Plots of the same select “Peaks” datasets in principal component space. Separability of clusters increases as the spread and width decrease. **C**. Heatmaps of the optimal results (based on the highest silhouette score) for each of the nine algorithms, including the recovered optimal number of clusters, the highest silhouette score, and corresponding adjusted rand index. **D**. Fidelity of recovering the ground truth number of clusters. Six is the ground truth number of clusters. Different algorithms are represented by different colors, as shown in the plot on the right measuring the deviation between the identified number of clusters (“BestK”) and the ground truth number of clusters (“GT”). **E**. The silhouette score, a measure of cluster separation, for the identified optimal number of clusters for each algorithm. **F**. The adjusted rand index, a measure of how correctly each neuron was assigned to its ground truth cluster, for each algorithm.

**Fig. S4.**
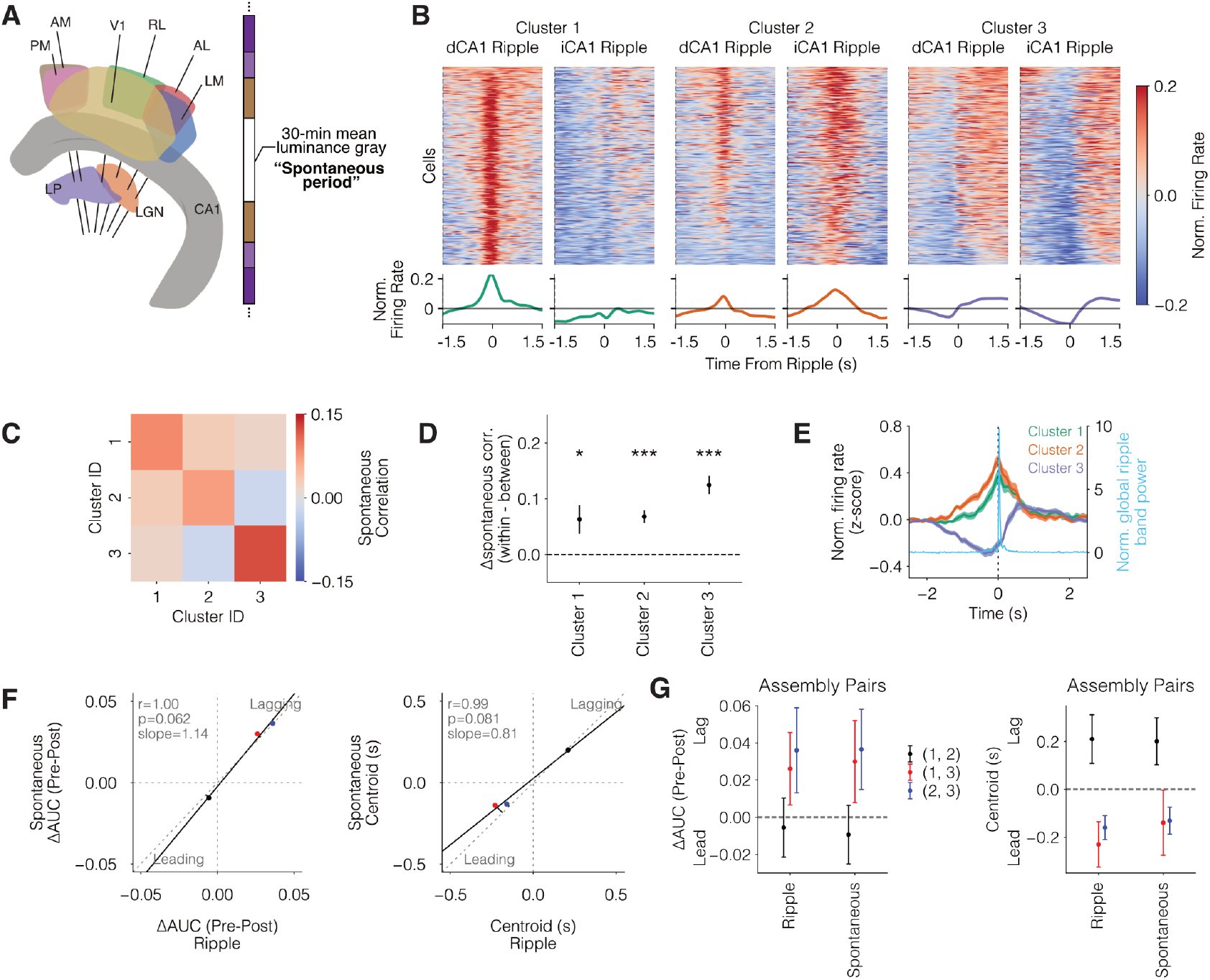
Identification using the clustering pipeline of sharp wave ripple-responsive visual cortex assemblies and their replayed sequences during and outside of ripple periods. **A**. Neuropixels probe locations and recording session timeline. Neuropixels probes targeted six visual cortical areas and also passed through the hippocampus and thalamus. Analysis used data from a thirty-minute spontaneous period during which mean luminance gray was presented. **B**. Ripple-responsive visual cortex neurons were clustered into three groups based on average activities around the two ripple types (see methods). The top row shows a heatmap of individual cell firing rates around each ripple type. The bottom row shows the average firing rate for each cluster around each ripple type. **C**. Heatmap showing spontaneous correlations between each pair of clusters based on their firing activity outside of ripple periods. **D**. The difference in within-cluster versus between-clusters spontaneous correlations outside of ripple periods. All clusters are assemblies. *p<0.05, **p<0.01, ***p<0.001. Error bars are s.e.m. **E**. Assembly activity sequence around global ripples. Error shading is s.e.m. **F**. ΔAUC and centroid time of the cross correlogram of pairwise assembly sequences is plotted for all cluster pairs during (ripple) and outside (spontaneous) of ripple periods (see methods). **G**. ΔAUC and centroid time of the cross correlogram of pairwise assembly sequences during ripple and spontaneous periods, showing maintenance of the sequence across both periods. Error bars are 95% confidence intervals estimated from a hierarchical bootstrap.

**Fig. S5.**
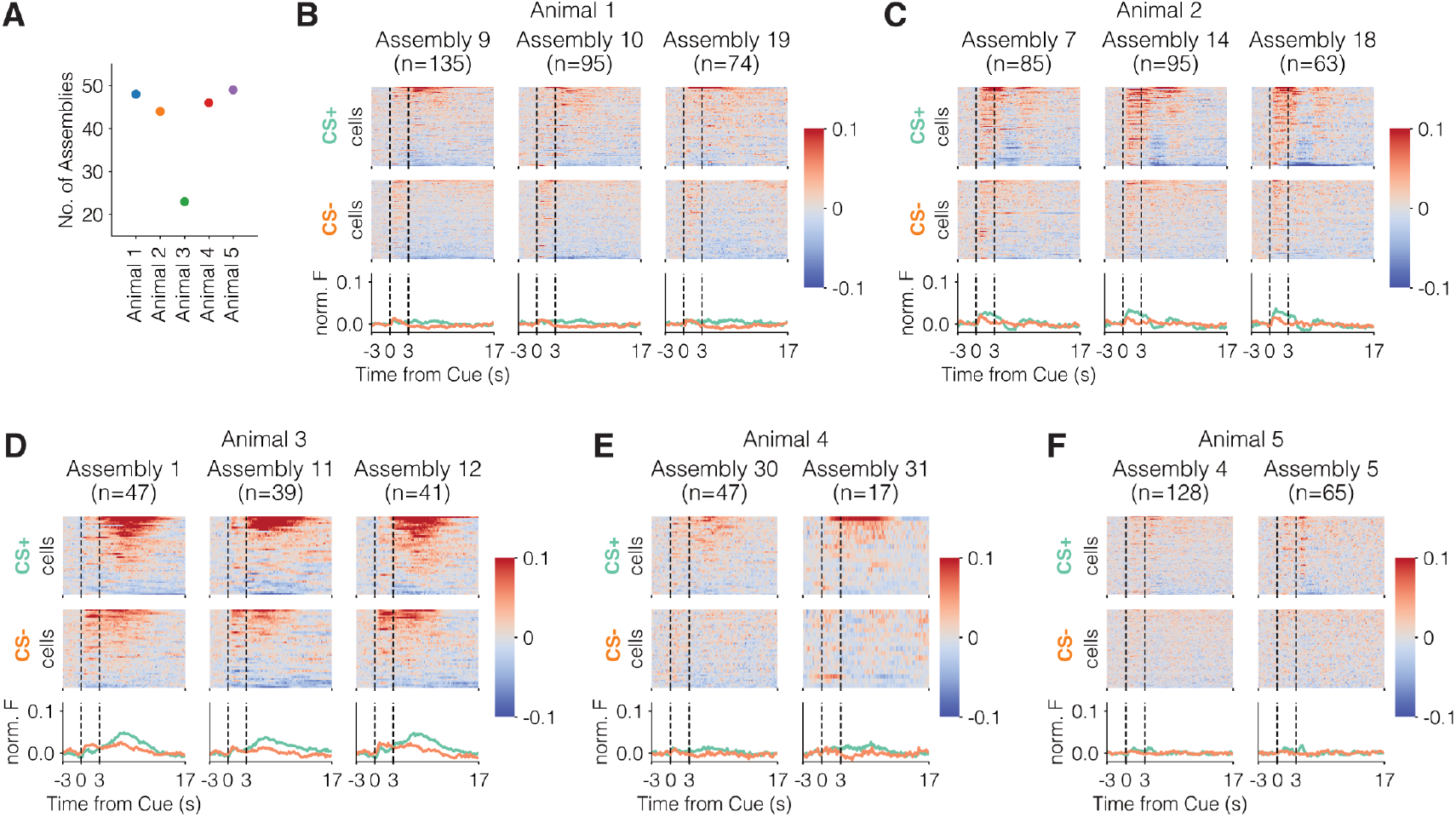
Results of a conventional assembly detection pipeline on OFC trained data from Lopes dos Santos et al. 2013 (61) showing that the task related activity of assemblies was highly redundant. **A**. Number of assemblies that were found for each subject animal using the method outlined by Lopes dos santos et al. 2013 (61). This method can only be utilized on data from an individual session of an individual animal. Many more assemblies were found compared to our pipeline, suggesting that some are redundant in terms of each assembly’s average activity profile. **B-E**. Examples of redundant assemblies with similar average cue-reward activity profiles for each animal.

**Fig. S6.**
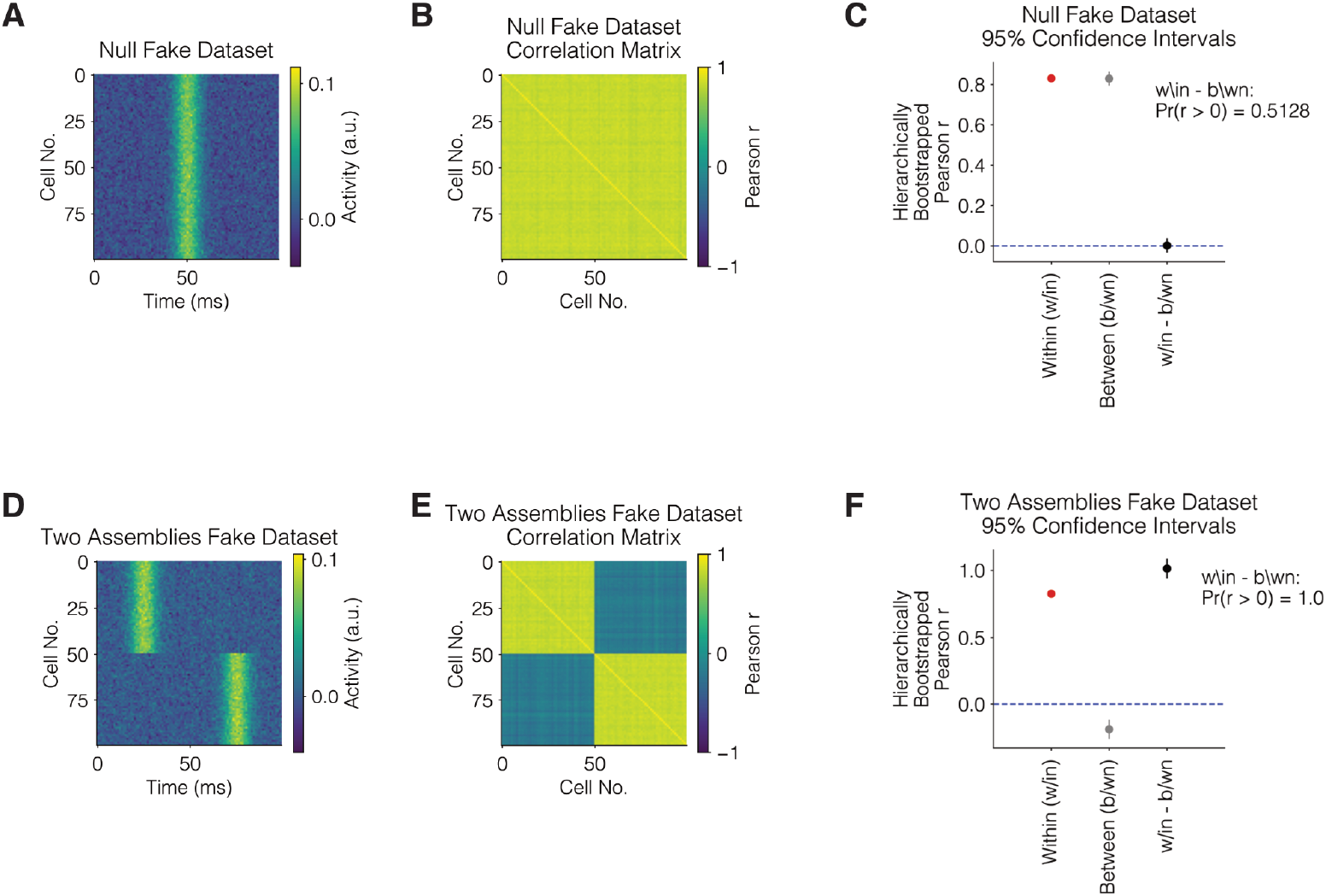
Validation of hierarchical bootstrapping of assembly detection using ground truth simulated data. **A**. Heatmap of a fake dataset we used to empirically check our method of hierarchically bootstrapping within cluster, between cluster, and within – between cluster Pearson r values. Cells 1-50 comprised cluster 1 and cells 51-100 comprised cluster 2. **B**. Correlation matrix of the null fake dataset of pairwise cell Pearson r correlations. All Pearson r values are approximately the same. **C**. Hierarchically bootstrapped 95% confidence intervals of Pearson r values. We empirically calculated the proportion of bootstrapped samples valued above 0, Pr(r > 0). **D**. Heatmap of a second fake dataset we used to empirically check our method of hierarchically bootstrapping within cluster, between cluster, and within – between cluster Pearson r values. Cells 1-50 comprised cluster 1 and cells 51-100 comprised cluster 2. **E**. Correlation matrix of the null fake dataset of pairwise cell Pearson r correlations. Pearson r values within a cluster are now much higher than between clusters. **F**. Hierarchically bootstrapped 95% confidence intervals of Pearson r values.

**Fig. S7.**
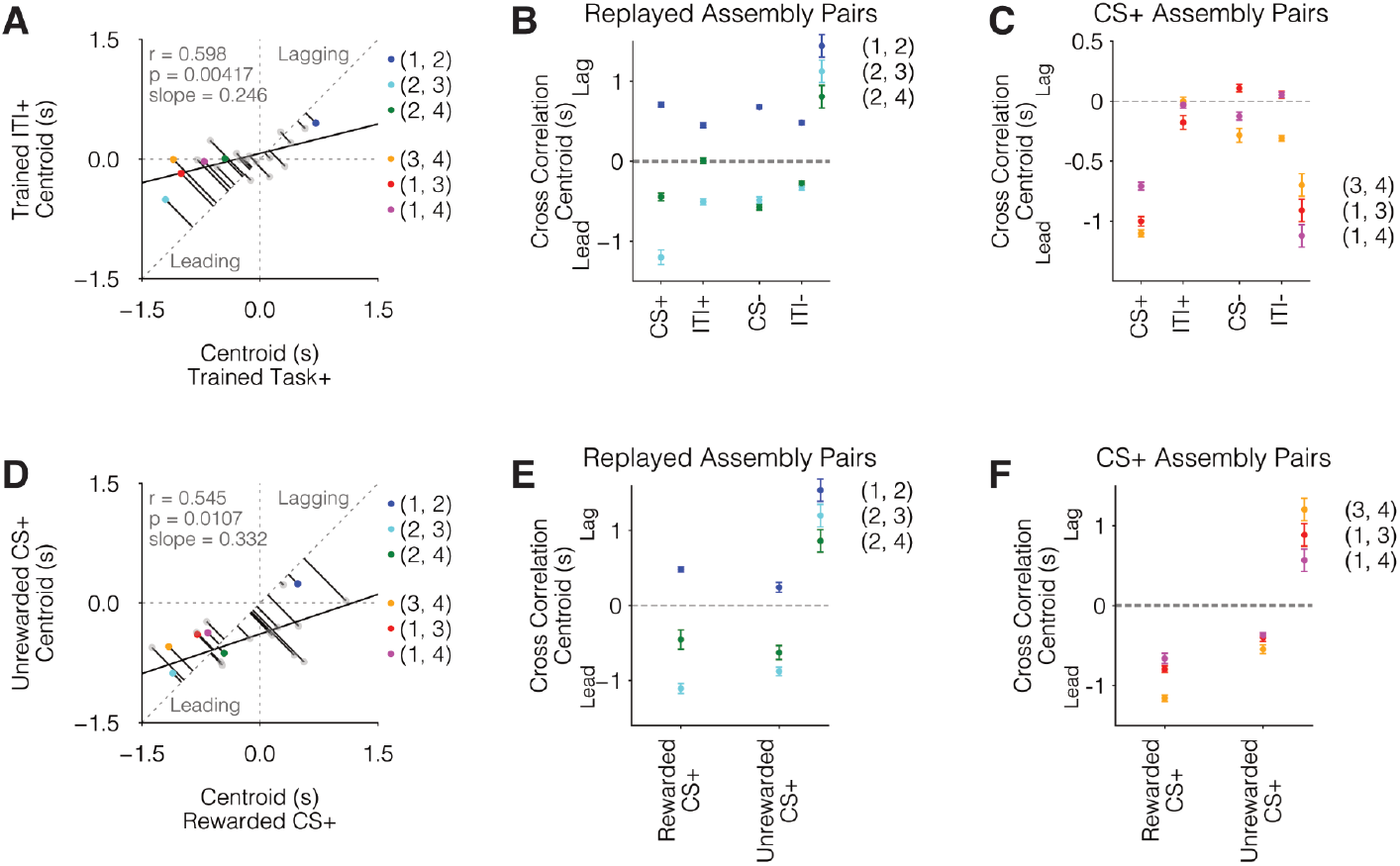
Cross correlogram centroid times of consolidation and retrieval assembly pairs during CS+, CS-, ITI+, and ITI-periods on the trained session, and CS+ unrewarded on the 50% reward probability session. Note that centroid times remain unchanged if the entire cross-correlogram is multiplicatively scaled. **A**. The sequential relationship between assemblies was measured using the area under the curve of the cross correlogram (CCG) (Fig 3) as well as the centroid of the CCG as shown here. Centroid time is plotted for all cluster pairs during CS+ periods (0-17 s following CS+ onset) and ITI+ periods (ITIs following CS+ trials). **B**. Centroid time of memory consolidation-like “replayed sequences” during CS+, ITI+, CS- and ITI-periods. Error bars are 95% confidence intervals estimated from a hierarchical bootstrap. **C**. Centroid time of memory retrieval-like “CS+ sequences” during CS+, ITI+, CS- and ITI-periods. Error bars are 95% confidence intervals estimated from a hierarchical bootstrap. **D**. Centroid time is plotted for all cluster pairs during CS+ rewarded vs unrewarded periods on the 50% reward probability session. **E**. Centroid time of “replayed sequences” during CS+ rewarded and unrewarded periods. Error bars are 95% confidence intervals estimated from a hierarchical bootstrap. **F**. Centroid time of “CS+ sequences” during CS+ rewarded and unrewarded periods. Error bars are 95% confidence intervals estimated from a hierarchical bootstrap.

**Fig. S8.**
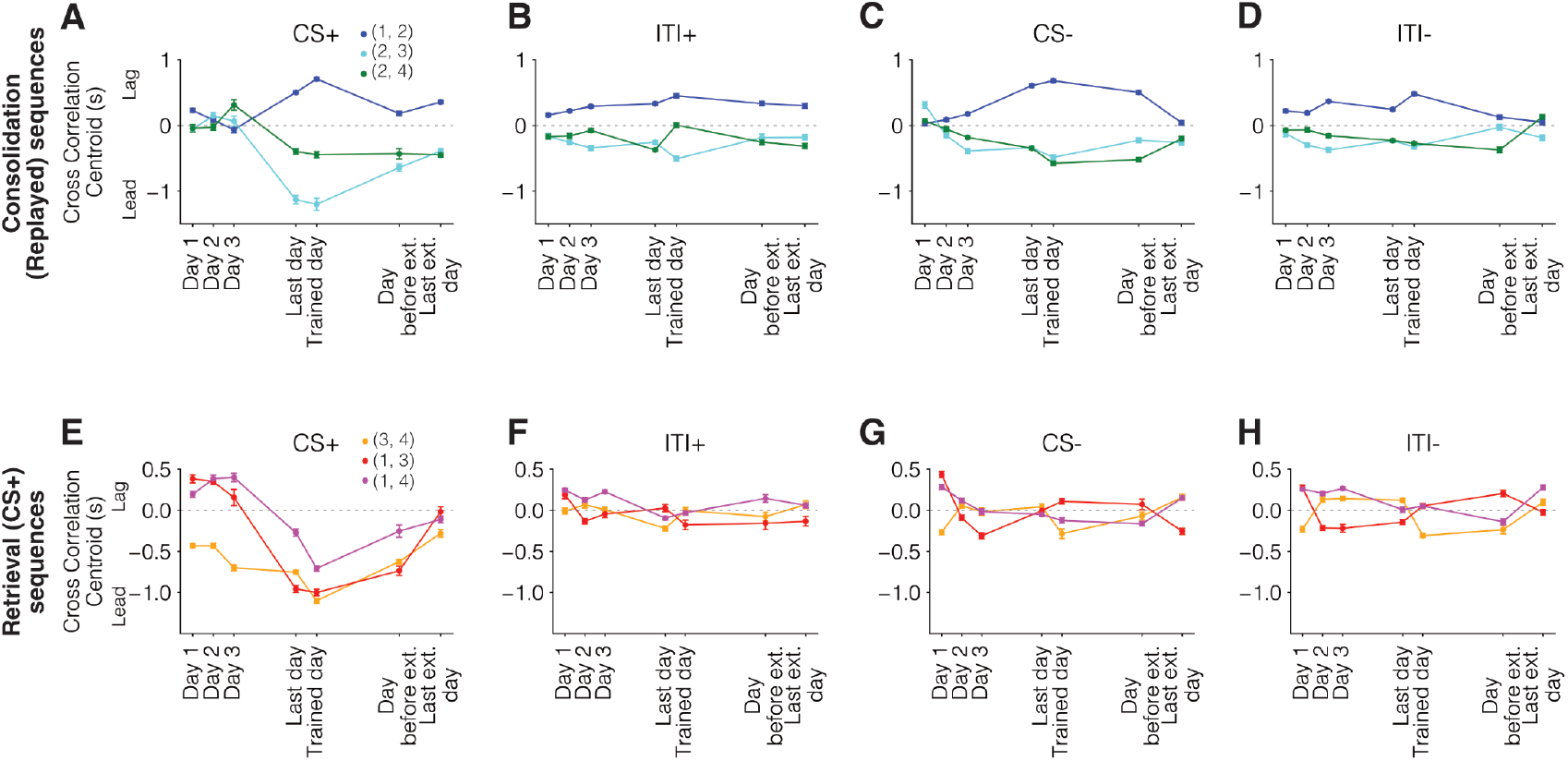
Cross correlogram centroid times for consolidation and retrieval assembly pairs during CS+, ITI+, CS-, ITI-throughout learning and extinction. **A-D**. Centroid time of consolidation “replayed sequences” across days of interest for all time periods. Error bars are 95% confidence intervals estimated from a hierarchical bootstrap. **E-H**. Centroid time of retrieval “CS+ sequences” across days of interest for all time periods. Error bars are 95% confidence intervals estimated from a hierarchical bootstrap.

**Fig. S9.**
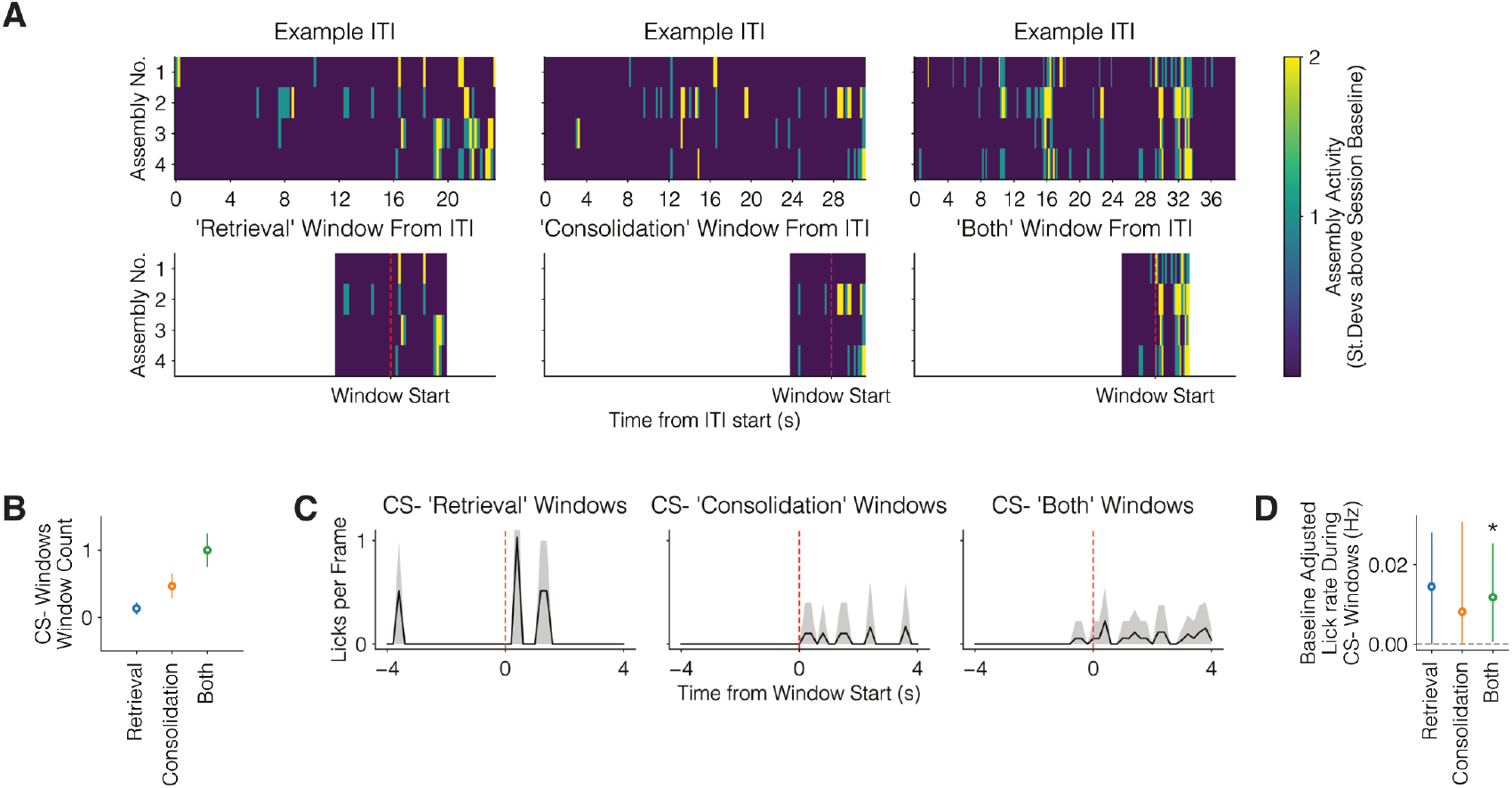
Replay window identification and CS- replay window results. **A**. Example of three different replay types. Three example inter-trial intervals (top row) along with examples of each of the three types of replay windows are shown (bottom row). We classified a window type depending on which assemblies had significantly high activity (2 standard deviations above session baseline) during a window. The same process was used to extract replay windows from CS-trial periods between [0,17s] from CS-cue onset. **B**. Number of identified ITI replays per session of either retrieval only sequences (CS+ sequences), single consolidation pairs (“consolidation” representing a single pair of replayed sequences (1,2), (2,3) or (2,4)), and multiple consolidation pairs, which include both consolidation and retrieval sequences (“both”). **C**. PETH of licking aligned to the above replay types. Error shading represents 95% confidence intervals from hierarchical bootstrapping. **D**. Baseline subtracted lick rates during the 4s after replay onset. Error bars represent 95% confidence intervals from hierarchical bootstrapping (*p<0.05, Bonferroni corrected).

## Supplementary table 1: Statistical Results

**Table 1.**
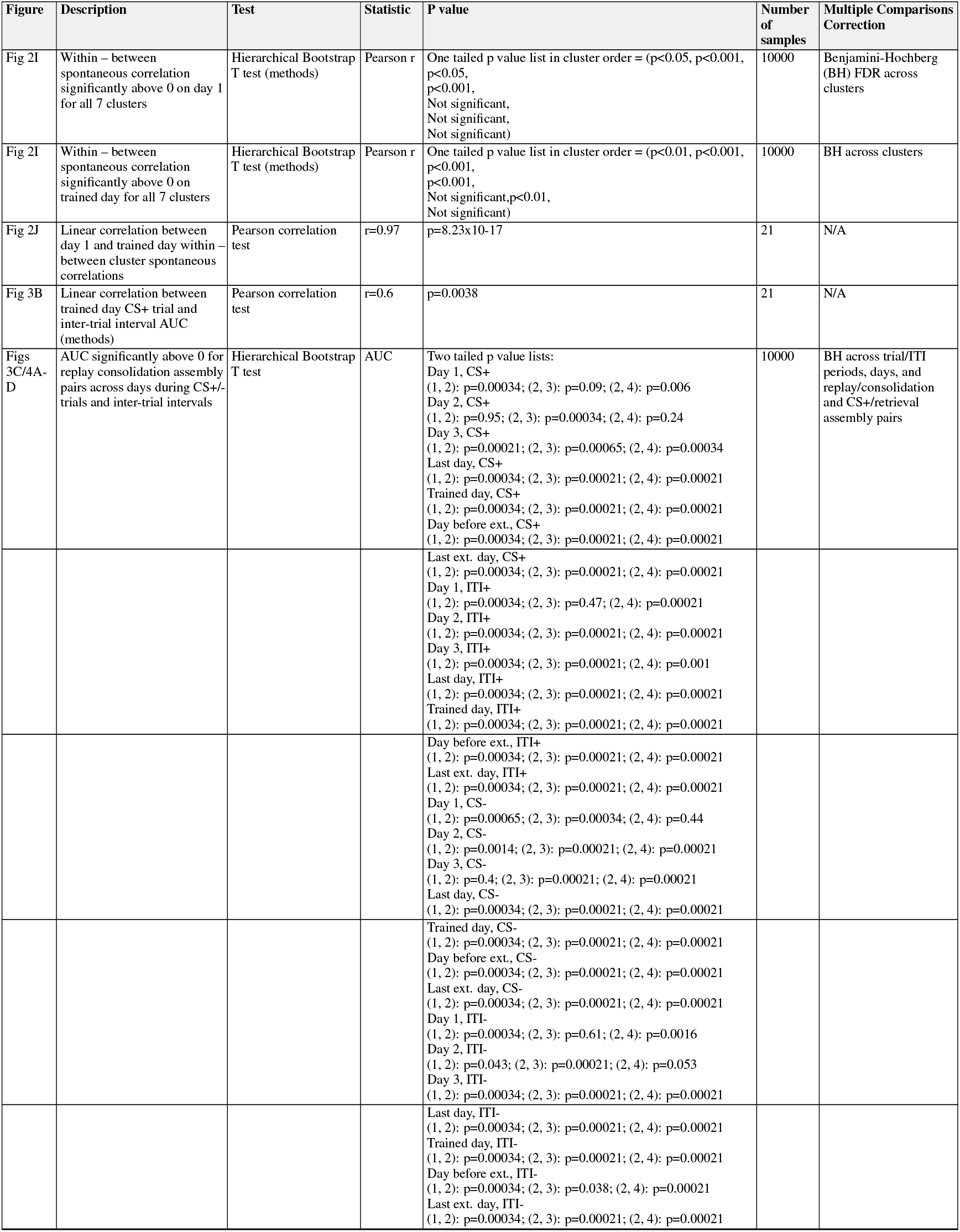
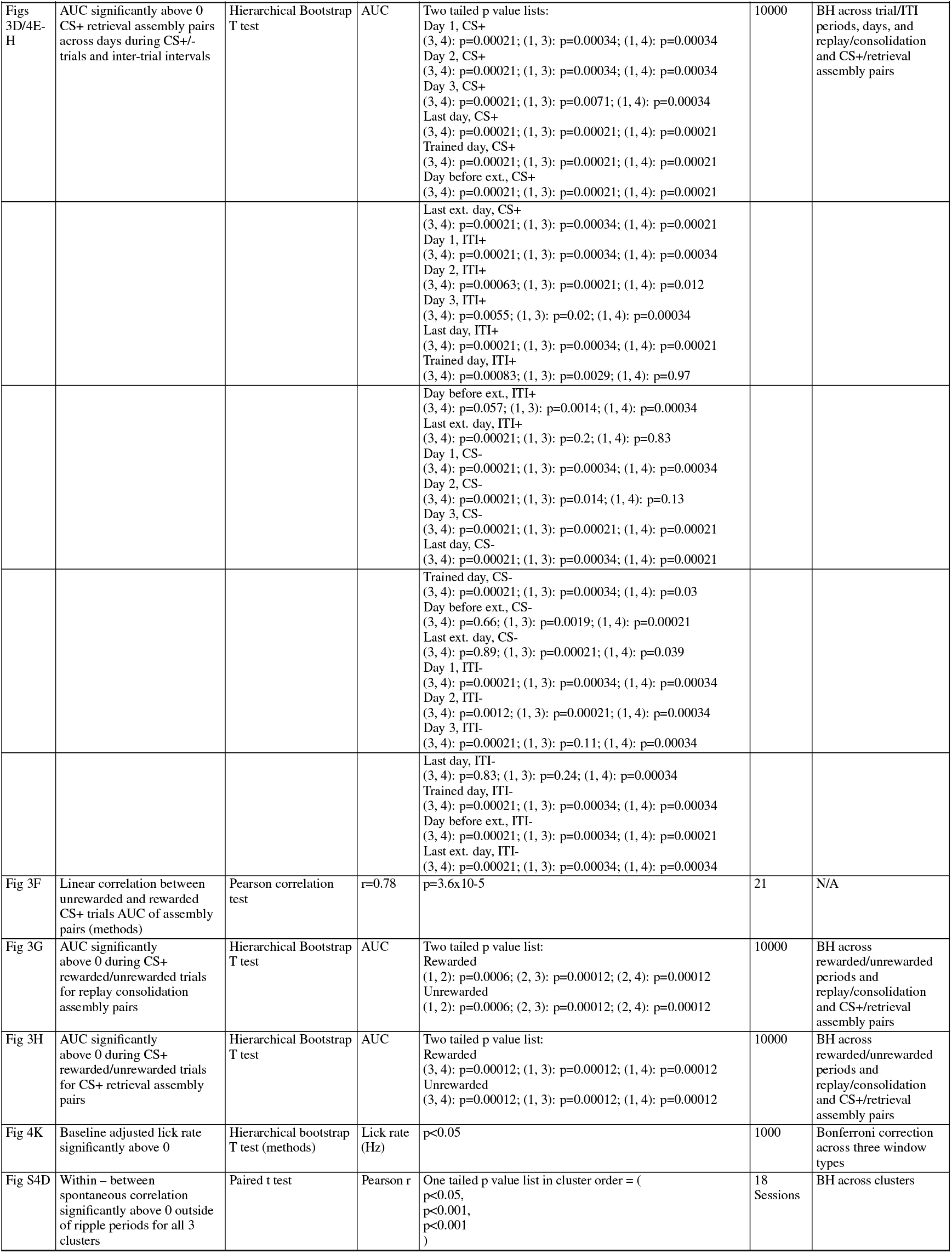
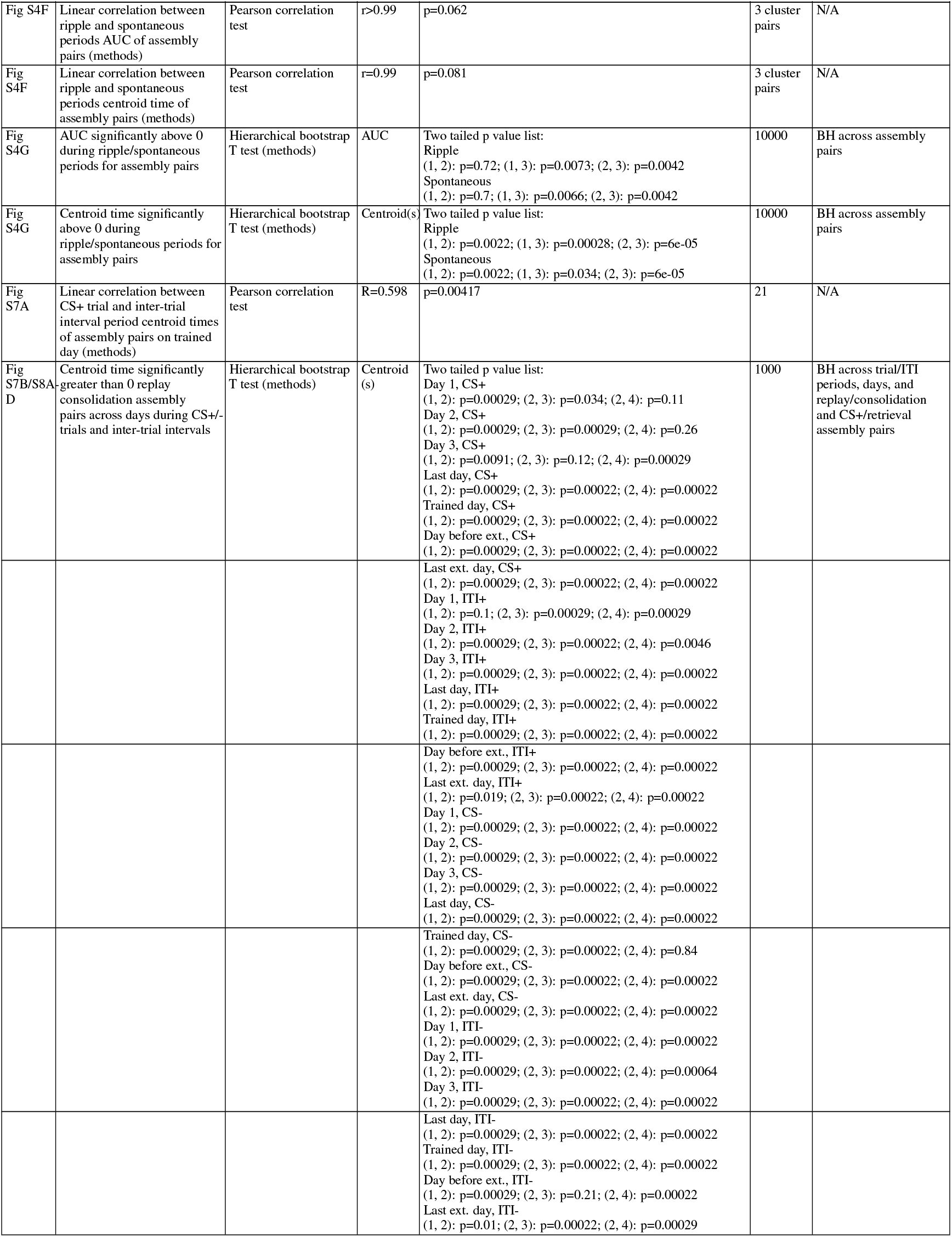
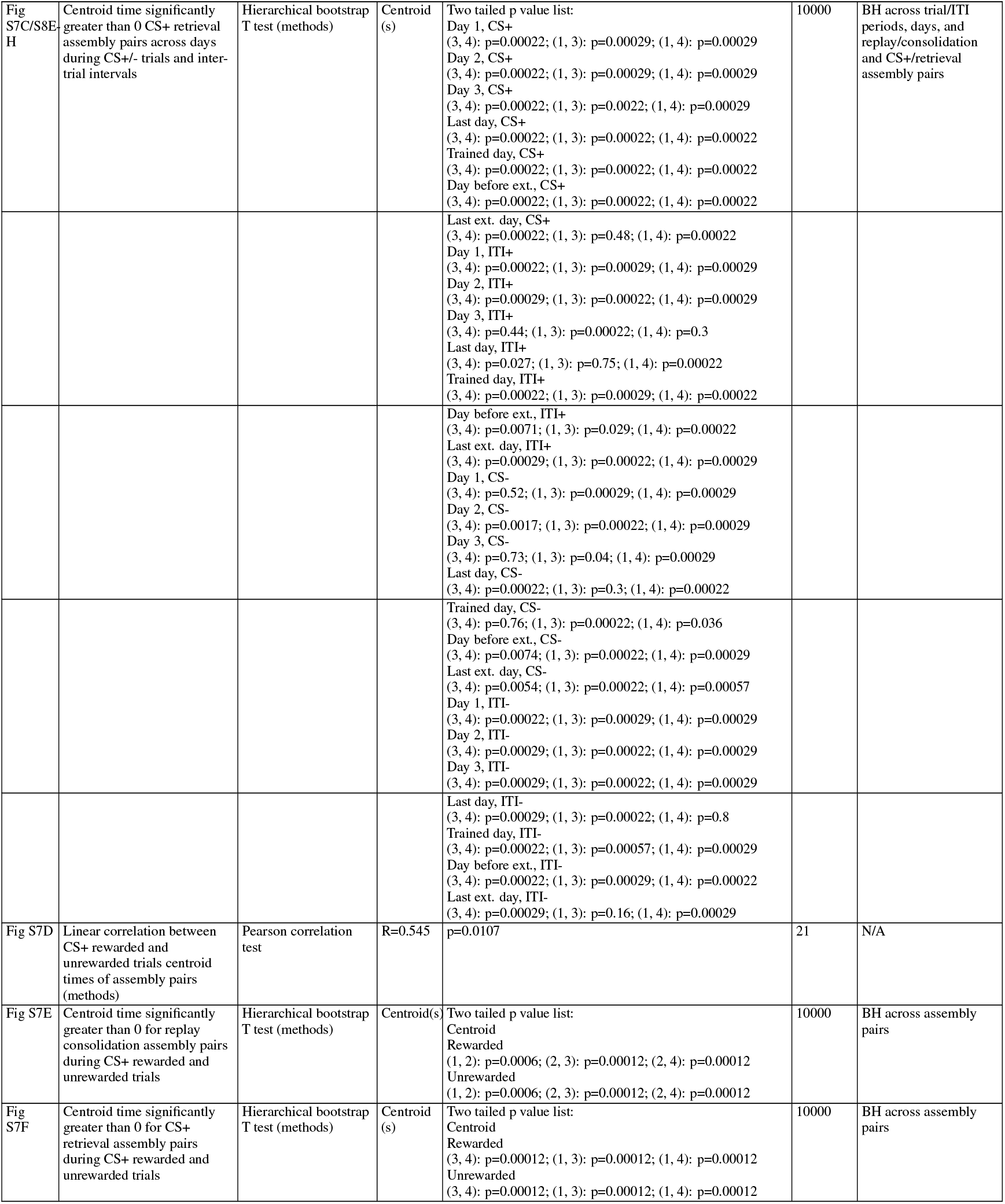
Table 1: Statistical results

## Methods

### Ground truth clustering simulations (Fig 1)

To evaluate which clustering algorithm to use for PETH separation in the assembly discovery pipeline, we generated three simulated ground truth datasets: one based on real data, and two hypothesized types of co-activity patterns. We refer to the first as the “Fake OFC” datasets (**Fig 1**) and the latter two as the “Shape” datasets (**Fig S2**) and “Peak” datasets (**Fig S3**).

To generate the “Fake OFC” dataset, we merged OFC neural clusters 7, 8, and 9 from previous data (55). We kept clusters 1 − 6 as is, for 7 total clusters. We calculated the sample mean and covariance matrices of the PSTHs calculated from cue aligned activity spanning [ − 3, 17] s from neurons in each cluster. We then generated several, entirely new, synthetic datasets using the python library NumPy *random*.*multivariate*_*normal* function. To generate a dataset, we scaled each covariance matrix per cluster by a coefficient alpha, parameterizing the separability of neurons in data space. Then we generated 100 new simulated “neurons” per cluster based on that cluster’s true sample mean in the data and the scaled covariance matrix. We sampled values of alpha between [0.05, 1] in increments of 0.05 to generate 20 datasets.

We generated the “Shape” simulation datasets to model neurons that share a common temporal shape of activity within each cluster, while different clusters differ in the number of distinct peaks in that shape. We simulated 5 clusters of 25 neurons each with 125 time bins for each neuron’s timeseries with a bin size of 0.5. For cluster k, each neuron’s time series was constructed from k Gaussian components (i.e., cluster 1 has one Gaussian, cluster 2 has two Gaussians, … cluster 5 has five Gaussians). The peak times were arranged as such: 1.5, 12.5, 25, 37.5, and 50. All Gaussian components shared the same standard deviation, and their peak times were arranged so that the first component’s peak time was identical across clusters (so cluster 2’s first peak aligns with cluster 1’s peak), then subsequent peaks were placed with adjacent offsets. To control separability, we scaled the amplitudes of all Gaussian components except the first by a common multiplier α, and generated datasets for α ranging [0, 0.66] in increments of 0.06. We added stationary Gaussian noise using NumPy’s random.normal function with default parameters and uniformly sampled noise NumPy’s random.uniform with range [0.01*M, 0.1*M] (M is the maximum value of a given time series) to each time series and normalized time series to sum to 1.

We constructed the “Peak” datasets to simulate neurons that have a Gaussian shaped curve with 6 clusters of simulated neurons with 20 neurons each. We used 200 time bins for each neuron’s time series with a bin size of 1. Neurons in each cluster had a shared peak time of activity. Each cluster had a unique peak time. We selected peak times of [50, 60, 95, 105, 140, 150], and parameterized data generation by mean noise, controlling the within cluster variability of single neuron peak times, and standard deviation, controlling the spread of individual neurons’ Gaussian shaped activities. As mean noise increases, cluster identity of neurons becomes harder to correctly recover, and when mean noise is greater than 5, peak times between adjacent clusters can overlap. We generated datasets with mean noise values between [0, 6] in increments of 1, and for each mean noise parameter value, we sampled standard deviation values of [0.6, 3.6] in increments of 0.6, then applied the exponential function to the sampled standard deviation value. We added stationary Gaussian and uniformly sampled noise as described above to each time series and normalized time series to sum to 1.

In **Fig S1**, we tested several algorithms (80%/90%/95% explained variance, scree plot elbow, Marchenko-Pastur cutoff, Kaiser rule, Broken Stick rule, Minka’s MLE rule, Optimal Hard Threshold, Horn’s parallel analysis) that automatically selected the number of principal components to retain after PCA in combination with several clustering algorithms and variants available in scikit learn (KMeans, Agglomerative across all linkage types, KMeans/Discrete/ClusterQR Spectral Clustering, DBSCAN, HDBSCAN, OPTICS) on these three simulated ground truth datasets. The entire pipeline was tested to identify the appropriate number of retained principal components and clustering algorithms and associated parameters that maximized performance metrics described later.

The approaches to identify number of principal components are described in detail below.

#### 80%/90%/95% explained variance

Choose the smallest number of principal components so that the cumulative proportion of variance explained exceeds a threshold (e.g., 80%, 90%, 95%).

#### Scree-plot “elbow”

Plot the eigenvalues (or variance explained) in descending order and look for the “elbow” point where the drop flattens out. Then retain all components before the elbow. This is a heuristic reflecting diminishing returns in variance explained beyond a point where additional components are likely due to noise in the data (100).

#### Marchenko–Pastur cutoff

Compute the theoretical upper bound of the bulk eigenvalue distribution under a null (noise) model and retains only components whose eigenvalues exceed that bound. Above the bound, components are treated as noise (101).

#### Kaiser rule (Kaiser-Guttman criterion)

Retain those components whose eigenvalue is equal to or greater than 1 (for correlation-matrix PCA) by the logic that each component should explain at least as much as one original variable (102).

#### Broken Stick rule

Compare the observed variances explained by each component to the expected variances from a “broken-stick” distribution, generated by randomly distributing total variance amongst components. Retain components whose observed variance exceeds the broken-stick expectation (103).

#### Thomas Minka’s Maximum Likelihood Estimation rule

Treats PCA as probabilistic/density estimation. First compute the Bayesian model evidence for different numbers of components, then select the k that maximizes evidence; effectively an automatic “maximum-likelihood” choice of dimension (104).

#### Optimal Hard Threshold for singular values

Derived in high-dimensional settings, this method sets a fixed threshold on singular-values above which components are retained. Below that threshold, components are considered noise (105). We used the optht python library optht.optht.

#### Horn’s Parallel Analysis

Create many random datasets (same size and variable count) of uncorrelated variables, compute their eigenvalues, and compare to observed eigenvalues. Then retain components whose eigenvalues exceed the corresponding percentile of the random-data eigenvalues (106). We used the horns python library function *horns*.*parallel*_*analysis*.

For clustering followed by the above PCA dimensionality reduction, we used the scikit-learn sklearn.cluster library to run and test all clustering algorithms listed below. Unless otherwise specified below, default parameters were used. We varied all n_cluster parameters from 1 to 10.

#### KMeans

Partitions the data into k clusters by repeatedly assigning points to the nearest cluster centroid and updating centroids to minimize the within-cluster sum of squared distances. The n_clusters parameter selects the number of clusters. The n_init selects the number of times to run with different initial seeds and was set to 1 (107).

#### Agglomerative/Hierarchical Clustering

Starts with each data point as its own cluster and successively merges the two closest clusters according to a chosen linkage criterion until a stopping condition. We tested each linkage type (‘ward’, ‘complete’, ‘average’, ‘single’), treating it like a parameter and selecting the best performing linkage type for each dataset. The n_clusters parameter selects the number of clusters. There is also the option to restrict the data points that get merged via the nearest neighbor’s connectivity graph. We did so for each linkage type, varying the number of nearest neighbors by a factor of 20 (108, 109).

#### Spectral Clustering

Constructs a similarity (affinity) graph of the data, computes an eigenvector embedding of the graph Laplacian, then clusters points in that embedding via KMeans, discretization, or cluster qr methods to capture non-convex structure (66, 110, 111). The n_clusters parameter selects the number of clusters. We varied the number of nearest neighbors used by a factor of 20 when computing the nearest neighbor’s graph for use as the affinity graph. We tested all three methods of assigning cluster points. We used the eigen_solver ‘arpack’.

#### DBSCAN

Finds clusters as regions of high point-density by grouping points that have at least a minimum number of neighbors within a radius *ϵ* and marks low-density points as noise. We systematically varied core parameters eps (maximum radius between neighbors) and min_samples (minimum number of points within that radius to be a “core” point) which control how dense a region must be to form a cluster and how many clusters and outliers result by factors of 10 (112, 113).

#### HDBSCAN

Builds on DBSCAN by constructing a hierarchical tree of density-based clusters over varying *ϵ* values and then selecting flat clustering based on cluster stability (114, 115). We varied the min_cluster_size (minimum size for any cluster) and min_samples (minimum number of neighbours for a core point, defaults to min_cluster_size if unspecified) parameters by factors of 10.

#### OPTICS

Orders points by reachability distance to reveal the clustering structure at multiple density levels and can extract clusters at varying densities (thus generalising DBSCAN’s fixed *ϵ* approach). We varied the parameters min_samples (minimum number of points in a neighbourhood for a core point), xi (steepness threshold controlling cluster boundary extraction) and min_cluster_size (minimum size for extracted clusters) which govern core-point selection, maximum radius, and extraction sensitivity by factors of 10 (116).

The reason we tested several criteria for number of retained principal components is that we observed that retaining too many PCs can result in over-clustering. We observed that the number of clusters is directly proportional to the number of PCs retained for a wide range. Therefore, retaining too many PCs causes the number of recovered clusters to go well beyond the ground truth number of clusters. Hence, selecting the correct number of PCs is crucial.

We also selected the number of nearest neighbors in a range without values close to 0 or number of samples. This is because these end ranges result in significantly lower performance metrics.

We measured three metrics for dimensionality reduction and clustering algorithm performance: BestK, Silhouette Score, and Adjusted Rand Index. BestK is the number of identified clusters (number of unique cluster labels). Silhouette Score is a commonly accepted intrinsic measure of cluster separation. We utilized the cosine distance metric for measuring silhouette score for all analyses. Adjusted Rand Index measures the similarity of clustering labels to the ground truth labels. We used the scikit-learn *sklearn*.*metrics*.*silhouette*_*score* and *sklearn*.*metrics*.*adjusted*_*rand*_*score* functions to compute them.

We observed that several dimensionality reduction methods (excluding the 80%/90%/95% empirical rules) similarly selected the correct number of PCs for downstream clustering, so we used the widely employed and computationally simple scree plot elbow method. For a given algorithm on a dataset, the optimal parameters were determined by the highest silhouette score over grid search. Then, the BestK and Adjusted Rand index values were calculated for those parameters. We measured overall performance for each clustering algorithm across the three datasets by calculating the area under the curve of each metric across dataset parameters.

### Electrophysiology data

We used an Allen Institute Neuropixels dataset (Allen Brain Observatory – Neuropixels Visual Coding) freely available to the public (https://allensdk.readthedocs.io/en/latest/visual_coding_neuropixels.html). Specifically, we used the ‘Functional Connectivity’ dataset of neural activity recorded from up to six Neuropixels probes while head-fixed mice were exposed to a variety of visual stimuli, including a 30-minute block of mean luminance gray screen. Analyses were conducted on this 30-minute block. We followed the same data preprocessing and exclusion criteria as previously published (68). In total, 20 mice (16 males) were used for the clustering, including 13 wild-type (WT), two Pvalb-IRES-Cre/wt;Ai32(RCL-ChR2(H134R) EYFP)/wt, three Sst-IRES-Cre/wt;Ai32(RCL-ChR2(H134R) EYFP)/wt, and two Vip-IRES-Cre/wt;Ai32(RCL-ChR2(H134R) EYFP)/wt mice. We also used a previously published pipeline for hippocampal sharp wave ripple detection (cite). Three types of ripples were identified: dorsal hippocampal ripple, intermediate hippocampal ripple, and global hippocampal ripple. Clustering was performed on PETHs of visual cortical neurons aligned to dorsal and intermediate ripples, but cross-correlation analysis during ripples was performed on global ripples. Details of clustering and cross-correlation analysis are described later.

### Behavior and imaging data (Fig 2-4)

All experimental procedures were approved by the Institutional Animal Care and Use Committee of the University of North Carolina and accorded with the Guide for the Care and Use of Laboratory Animals (National Institutes of Health). These data were collected as described previously (55). Briefly, adult male wild type C57BL/6J mice (Jackson Laboratories, 6-8 weeks, 20-30 g) were group housed with littermates and acclimatized to the animal housing facility until surgery. Survival surgeries were stereotaxically performed while maintaining sterility. Following recovery, animals used for behavioral studies were water deprived to reach 85-90% of their pre-deprivation weight and maintained in a state of water deprivation for the duration of the behavioral experiments.

Water deprived mice were first trained to lick for random unpredictable sucrose (10-12.5%, 2.5 µL) deliveries in a conditioning chamber. On subsequent days, mice received one of two possible auditory tones (3 kHz pulsing tone or 12 kHz constant tone, 75-80 dB) that lasted for 2 seconds. A second after the cues turned off, the mice received a sucrose reward following one of the tones (CS+), while the other tone resulted in no reward (CS-). The identity of the tones was counterbalanced across mice. A total of 50 CS+ and CS-each were presented in a pseudorandom order. The intertrial interval between two consecutive presentations of the cues was drawn from a truncated exponential distribution with mean of 30 s and a maximum of 90 s, with an additional 6 s constant delay. Animals were considered trained after the area under a receiver operating curve between anticipatory licks on CS+ vs CS-trials became larger than 0.7 on at least 2 consecutive sessions or larger than 0.85. Two contingency degradation experiments were performed as described previously (55), with reward probability reduced 50% in one (**Fig 3E**) and background unpredictable rewards introduced in the intertrial interval in the other while CS+ fully predicted reward. After retraining with full contingency, the CS+-reward pairing was extinguished for at least two days.

GCaMP6s was virally expressed under a CaMKIIα promoter to image single-cell calcium changes using 2-photon microscopy through a gradient refractive index (GRIN) lens implanted in the ventral/medial OFC (55). Data were acquired with a 6 frame online averaging at an effective frame rate of 5 Hz. Motion correction and cell extraction were performed as explained previously (55). Imaging acquisition was triggered by a custom Arduino code and a TTL output of every frame was sent as an input to the Arduino. The imaging acquisition was triggered off at the end of the behavioral session. In every mouse, one z-plane was imaged throughout acquisition so that the same cells could be tracked through learning. Only such longitudinally tracked neurons are included here.

### Data Analysis

All statistical tests involving multiple comparisons utilized the Benjamini-Hochberg correction. P values were directly estimated using hierarchical bootstrap.

#### Identifying task responsive OFC neurons

We used a Gaussian family GLM with an identity link function to filter out neurons that were inactive during the task. Continuous fluorescence was modeled under a normality assumption for simplicity. Each neuron’s median normalized fluorescence [-3,17] s from cue onset was selected and concatenated. Median normalization was done as (F(t)-median(F(t)))/(max(F(t))-min(F(t)) such that normalized fluorescence was close to 0 except during calcium events. The design matrix included an indicator variable and an intercept term. The indicator represented CS+ cues with a 1 and CS-trials with a 0 at each timepoint throughout the entire window. Parameters were estimated by maximum likelihood via iteratively reweighted least squares using the statsmodels python library GLM function, and parameter inference used t-tests with two-sided p-value. We identified active OFC neurons by keeping neurons whose indicator variable parameter had an associated p value less than 0.05. The GLM equation for a single neuron is as shown as below:

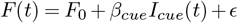

Where F(t) represents median normalized fluorescence on frame t, *β* represents the coefficients, I represents the indicator variable, and *ϵ* represents the error term. A total of 925 task responsive longitudinally tracked OFC neurons were used for analysis.

#### Clustering of OFC neurons

We clustered all OFC neurons pooled across animals that were recorded on the trained day (i.e., not just the longitudinally tracked neurons analyzed hereafter) by their median normalized fluorescence after filtering task inactive neurons using the above GLM. After filtering, the averaged fluorescence during the [-3,17] second window relative to cue onset was calculated separately for CS+ (rewarded) and CS-(unrewarded) trial types. The average fluorescence timeseries for each trial type was concatenated and z-normalized, comprising a 200-column vector of datapoints for each neuron.

As determined by our analysis in **Fig 1**, we applied principal component analysis for dimensionality reduction and kept the number of principal components determined by the scree plot elbow. We subsequently clustered neurons using the discrete spectral clustering algorithm variant using the scikit-learn function *sklearn*.*cluster*.*SpectralClustering* with the affinity matrix calculated using a k-nearest neighbor connectivity matrix. We tested the following numbers of nearest neighbors based on earlier ground truth simulations: 100, 200, 300, 500, 750, 1000, 1500, 2000, 2500. The number of clusters was also systematically varied from 2 to 20. The best combination of parameters was chosen by maximizing the silhouette score over a grid search of parameters. Each combination of parameters was run 100 times with a different random seed due to the stochasticity of the discrete spectral clustering algorithm. All subsequent analyses were repeated across the 100 iterations of cluster labels that were generated for the optimal parameters.

#### Clustering of Visual cortex neurons

The averaged firing rates of visual cortical neurons during the [-1.5, 1.5] second window relative to ripple onset were calculated separately for dCA1 and iCA1 ripple types. The averaged firing rate timeseries for each ripple type were concatenated and z-normalized for each neuron, comprising a single dataset. We followed the same dimensionality reduction and cluster procedure as above. Labels were shown to be stable across random iterations.

#### Correlation and Cross Correlation Analysis

To estimate the lag 0 correlation during task periods and inter trial intervals within and between assemblies, we calculated all pairwise correlations between neurons using the deconvolved fluorescence-inferred spiking activity during all time windows for each animal. Time windows for task periods spanned [0,17] s from cue onset, and time windows for inter trial intervals spanned 17 seconds after cue onset to the time of next cue onset. Inter trial intervals that were shorter than three frames were excluded. We then employed the hierarchical bootstrap sampling method directly on these pairwise correlations to get an estimate for the true lag 0 correlations for a given assembly. For each assembly, we hierarchically sampled animals and then neuron pairs for 10,000 bootstrapped samples. To estimate the within-assembly correlation, we sampled all pairs of neurons where both neurons belonged to a given assembly. To estimate between-assembly correlations, we sampled neuron pairs where one neuron belonged to an assembly, and the other neuron did not. For computing between-assembly lag 0 correlations for a specific assembly pair, we sampled neuron pairs where one neuron belonged to the first assembly, and the other neuron belonged to the second. Phipson-Smyth corrected p values for within cluster correlation minus between cluster correlation were directly estimated from hierarchical bootstrap samples (117).

To estimate cross correlations, the area under the curve (AUC), and centroid time derived from the cross-correlation curves between pairs of assemblies, we averaged the deconvolved activity within a single time window of all neurons grouped in an assembly and then computed the cross correlation between pairs of assemblies’ average activity with up to 4 seconds of lag on individual time windows. We used the same time windows as above for task periods and ITI periods. ITI periods shorter than 4 seconds were excluded from the analysis. Then we computed the cross-correlation AUC and centroid time for each time window. We z-normalized each cluster time series then computed the cross correlogram with the NumPy function correlate in default mode ‘valid’. Next, we hierarchically sampled animals and then time windows to come up with a bootstrapped estimate of AUC or centroid time for each assembly pair. We generated 10,000 bootstrapped samples to estimate means and 95% confidence intervals. 95% confidence intervals were calculated as the 2.5 and 97.5 percentiles of the bootstrapped distribution.

Lag 0 and cross correlation analyses were repeated using the same hierarchical bootstrap procedure for visual cortex neurons from the Functional Connectivity dataset. We calculated Lag 0 correlations within and between pairs of visual cortex cluster single neurons using activity during all time windows outside of any [-1.5,1.5] s window surrounding ripples. We term these windows outside of ripple windows as spontaneous windows. Adjacent ripples with overlapping windows were joined together. Cross correlations were computed using 4 seconds of lag during ripple windows and spontaneous windows. Phipson-Smyth corrected p values for within cluster correlation minus between cluster correlation were directly estimated from hierarchical bootstrap samples (117).

#### Replay window identification

We identified specific windows of time during inter-trial intervals (ITIs) and CS-trials where consolidation sequences ((lead, lag) assembly pairs (1,2), (2,3), or (2,4) are active), retrieval sequences (any combination of assemblies 1,3, and 4 are active with assemblies 1 and 3 being possible leads), or a combination of both (assemblies 1,2, and 3 or 1,2,3, and 4 are active with 1,2, and 3 being possible leads) were replayed. Replay windows were identified using the criteria listed below. These were developed to ensure that each replay was not contaminated by possible prior replays/assembly activations. Because many assemblies showed long periods of sustained activity (e.g., cluster 1 activation lasts longer than 6s during CS+ unrewarded trials), we required a baseline period of 6s without considerable assembly activation. All analyses were performed on deconvolved fluorescence traces. Standard deviation for assembly activity events was calculated based on an assembly’s deconvolved fluorescence activity for a whole session.

1. We identify potential replay windows when a leading assembly has a 2 standard deviation activity event. The replay window starts when this same assembly has a preceding 1 standard deviation event within 1 second. If a separate 1 standard deviation activity event was not detected, the window was set to start 3 frames (0.6 seconds) before the leading assembly 2 standard deviation event.
2. There must be at least one lead assembly 2 standard deviation activity event and at least one lag assembly 2 standard deviation activity event within 4 seconds of each other during a replay window.
3. Prior to window start, there must not be any 2 standard deviation activity events by any of the four assemblies in the 6 preceding seconds. There must also not be more than five 1 standard deviation activity events by any assembly.

If the criteria were met, a 4 second replay window was extracted following the window start along with a 4 second baseline period prior to window start. Replay windows from ITIs and CS-trials were identified using all the trials pooled from the last day of learning, trained day, and the day before extinction (all of which had 100% predicted reward probability following CS+). We then extracted each animal’s licking activity during those replay windows to hierarchical bootstrap 95% confidence intervals and Phipson-Smyth corrected p values across days and animals for licking PSTHs and baseline adjusted lick rates (117).

